# The genetic architecture of helminth-specific immune responses in a wild population of Soay sheep (*Ovis aries*)

**DOI:** 10.1101/628271

**Authors:** A. M. Sparks, K. Watt, R. Sinclair, J. G. Pilkington, J. M. Pemberton, T. N. McNeilly, D. H. Nussey, S. E. Johnston

## Abstract

Host-parasite interactions are powerful drivers of evolutionary and ecological dynamics in natural populations. Variation in immune responses to infection is likely to shape the outcome of these interactions, with important consequences for the fitness of both host and parasite. However, little is known about how genetic variation contributes to variation in immune responses under natural conditions. Here, we examine the genetic architecture of variation in immune traits in the Soay sheep of St Kilda, an unmanaged population of sheep infected with strongyle gastrointestinal nematodes. We assayed IgA, IgE and IgG antibodies against the prevalent nematode *Teladorsagia circumcincta* in the blood plasma of > 3,000 sheep collected over 26 years. Antibody levels were significantly heritable, ranging from 0.21 to 0.39 in lambs and from 0.23 to 0.57 in adults. IgA levels were strongly associated with a region on chromosome 24 explaining 21.1% and 24.5% of heritable variation in lambs and adults, respectively; this region was adjacent to two candidate loci, the Class II Major Histocompatibility Complex Transactivator (*CIITA*) and C-Type Lectin Domain Containing 16A (*CLEC16A*). Lamb IgA levels were also associated with the immunoglobulin heavy constant loci (*IGH*) complex on chromosome 18. Adult IgE levels and lamb IgG levels were associated with the major histocompatibility complex (MHC) on chromosome 20. This study provides evidence of high heritability of a complex immunological trait under natural conditions and provides the first evidence from a genome-wide study that large effect genes located outside the MHC region exist for immune traits in the wild.

**Author summary:** Host-parasite interactions are powerful drivers of evolutionary and ecological dynamics in natural populations. Variation in immune responses to infection shapes the outcome of these interactions, with important consequences for the ability of the host and parasite to survive and reproduce. However, little is known about how much genes contribute to variation in immune responses under natural conditions. Our study investigates the genetic architecture of variation in three antibody types, IgA, IgE and IgG in a wild population of Soay sheep on the St Kilda archipelago in North-West Scotland. Using data collected over 26 years, we show that antibody levels have a heritable basis in lambs and adults and are stable over lifetime of individuals. We also identify several genomic regions with large effects on immune responses. Our study offers the first insights into the genetic control of immunity in a wild population, which is essential to understand how immune profiles vary in challenging natural conditions and how natural selection maintains genetic variation in complex immune traits.

## Introduction

Host-parasite interactions are powerful drivers of evolutionary and ecological dynamics in natural populations. Variation in immune responses to infection is likely to shape the outcome of these interactions, with important consequences for the fitness of both host and parasite. In the wild, individuals are exposed to a range of micro- and macro-parasites as well as variable and challenging environments, leading to considerable variation in immune phenotypes in comparison to those in controlled environments (1–3). This has the potential to alter the effect of underlying genetic variation and importance of particular genes for resistance to infection, yet little is known about the specific genetic mechanisms driving immune responses under natural conditions. By investigating the genetic architecture of immune traits in the wild, we can determine the relative contribution of genetic, environmental and individual variation to identify the evolutionary forces shaping individual differences in immunity.

Studies in humans and livestock have shown that variation in immune traits is often heritable; that is, a significant proportion of phenotypic variance can be attributed to additive genetic effects (4–11). Genome-wide association studies (GWAS) in these systems have identified a number of genes of relatively large effect contributing to heritable variation, most notably the major histocompatibility complex (MHC) and cytokine genes (12–17). In wild populations, studies have investigated the heritability of immune traits, most often in birds (18–23), with candidate gene approaches further implicating MHC and cytokine regions in cases where significant associations are observed (23–28). However, these studies often focus on broad, non-specific immune phenotypes such as the phytohaemagglutinin (PHA) response, haematocrit levels and/or parasite burden, rather than specific immune responses to ecologically-relevant parasites (29–31). In addition, candidate gene studies focus on a small proportion of the genome and may fail to identify previously undiscovered coding or regulatory regions associated with immune trait variation (32,33). To our knowledge, there are no genome-wide association studies of specific immune phenotypes in the wild.

Domestic sheep (*Ovis aries*) and their gastrointestinal strongyle nematodes represent a well-understood host-parasite system, due to their agricultural and economic importance, with much recent interest in determining the genes underlying host resistance to these parasites (34). Of the strongyle parasites, *Teladorsagia circumcincta* is of major economic importance for domestic sheep in temperate regions (35) and has a simple direct life-cycle, with an infective L3 stage which develops to L4 stage within the gastric glands before emerging as sexually mature adult parasites which reside in the abomasum. Defence against *T. circumcincta* in lambs is associated with parasite-specific IgA antibody responses directed at worm growth and subsequent female fecundity (6,36), while in older animals a hypersensitive response, involving IgE antibodies, results in expulsion of incoming larvae from the mucosa (35). Anti-*T. circumcincta* IgA levels are moderately heritable in lambs and adults (6,37,38). Candidate gene and genome-wide studies have identified regions associated with FEC or protective immunological traits related to gastrointestinal nematodes, with candidate gene studies primarily focussed on interferon gamma (*IFNγ*) and the MHC (34). However, due to the focus on identifying individuals for selective breeding and the greater impact of parasite infections in lambs, most studies focus only on lambs, with only a few studies of adult ewes (38–41). As a consequence, we know relatively little about age-dependent genetic effects; indeed, differences in resistance loci between lambs and adults suggest that the genetic control of these mechanisms may differ (40). In both agricultural and wild systems, adult females contribute to pasture larval counts during the periparturient rise; therefore, understanding the genetic basis of resistance in young and old animals will improve our understanding of host-parasite dynamics and interactions in these systems (42).

The long-term individual-based study of the Soay sheep of St Kilda provides a powerful opportunity to understand the genetic architecture of immune traits at different ages under natural conditions. Soays are infected with several gastrointestinal strongyle nematodes common to domestic sheep, predominantly *Teladorsagia circumcincta, Trichostrongylus axei* and *Trichostrongylus vitrinus* (43,44). Strongyle nematode burden, in combination with harsh winter weather and low food availability, is a strong selective force on the sheep (43–46). Parasite-specific antibody responses are moderately heritable (26,47) and parasite-specific IgA levels and parasite-specific pan-isotype antibody levels have been shown to be negatively associated with strongyle faecal egg count (24,47). Recent examination of anti-nematode antibody isotypes (namely IgA, IgE and IgG) over a 26-year period showed that levels of IgG are positively associated with adult survival, and negative associations are observed between antibodies and FEC for all isotypes in lambs, but only for IgG in adults (48–50).

Previous examination of the genetic architecture of immune traits in Soay sheep using QTL mapping and candidate gene approaches failed to identify loci associated with parasite egg counts and pan-isotype antibody levels (26,51); however, a microsatellite polymorphism at the IFNγ locus in lambs had previously been associated with reduced faecal egg counts and increased parasite-specific IgA levels (24). Today, the majority of study individuals have been genotyped on the Illumina 50K OvineSNP50 BeadChip, and genome-wide association studies have identified genomic regions associated with traits such as horn morphology, body size and recombination rate (52–54). Here, we investigate the heritability and conduct genome-wide association studies of anti-*T. circumcincta* IgA, IgE and IgG levels from plasma samples collected from Soay sheep over a 26-year period. We show that antibodies are heritable and temporally stable over an individual’s lifetime, and that several genomic regions explain heritable variation in both lambs and adults.

## Methods

### Study population

The Soay sheep is a primitive breed of domestic sheep that was isolated on the island of Soay in the remote St Kilda archipelago several millennia ago, and has been living in unmanaged conditions since then (55). In 1932, >100 Soay sheep were moved to the larger island of Hirta after the evacuation of all human residents. The population now fluctuates between from 600 to 2,200 individuals. Approximately a third of the Hirta population lives in the Village Bay area, and these individuals have been the subject of a long-term study since 1985 (55). In April, around 95% of all lambs born in the study area are caught each year and individually tagged. Each August, as many sheep as possible from the study population are re-captured using temporary traps (55). At capture, whole blood samples are collected into heparin tubes, centrifuged at 3000 r.p.m. for 10 minutes, and plasma removed and stored at −20°C.

### Quantifying antibody levels

This study quantified antibody levels in animals that were caught and blood sampled in August between 1990 and 2015, comprising 6543 samples from 3190 individuals. Five samples from late-born lambs caught in August within 50 days of birth were excluded from the dataset, due to the potential presence of maternal antibodies and differences in development stage to other lambs. Levels of the antibodies IgA, IgG and IgE against antigens of the third larval stage of *Teladorsagia circumcincta* were measured using direct (IgA, IgG) and indirect (IgE) ELISAs. We used *T. circumcincta* L3 somatic antigen, provided by the Moredun Research Institute, as the capture antigen for all three assays diluted to 2µg/ml in 0.06M carbonate buffer at pH 9.6. 50µl of the diluted capture antigen was added to each well of a Nunc-immuno 96-microwell plate, which was covered and incubated at 4°C overnight. After washing the wells three times in Tris-buffered saline-Tween (TBST) using a plate washer, 50µl of the Soay sheep plasma sample diluted to 1:50 for IgA and IgE, and 1:12800 for IgG was added to each well. The plates were then covered and incubated at 37°C for 1 hour. Plates were then washed five times with TBST and 50µl per well of rabbit polyclonal anti-sheep IgA detection antibody conjugated to horseradish peroxidase (HRP) (AbD Serotec AHP949P) diluted 1:16000 was added to the anti-*T. circumcincta* IgA assay and 50µl per well of rabbit polyclonal anti-sheep IgG detection antibody conjugated to HRP (AbD Serotec 5184-2104) diluted 1:16000 was added to the anti-*T. circumcincta* IgG assay. For the anti-*T. circumcincta* IgE assay, 50µl per well of anti-sheep IgE (mouse monoclonal IgG1, clone 2F1, provided by the Moredun Research Institute) diluted 1:100 was added, followed by 1-hour incubation at 37°C, five washes with TBST and then 50µl per well of goat polyclonal anti-mouse IgG1-HRP detection antibody (AbD Serotec STAR132P) was added diluted to 1:8000 in TBST. All plates were then incubated at 37°C for 1 hour. Plates were then washed five times with TBST and 100µl of SureBlue TMB 1-Component microwell peroxidase substrate (KPL) was added per well and left to incubate for 5 minutes in the dark at 37°C. Reactions were stopped by adding 100µl per well of 1M hydrochloric acid and optical densities (OD) were read immediately at 450nm using a Thermo Scientific GO Spectrophotometer.

All results were measured as OD values due to the lack of standard solutions. To minimise confounding of capture year and age effects with plate to plate variation, each plate included samples from two years paired at random with different age groups on each plate. All plates were run in duplicate and duplicate sample ODs were removed if the coefficient of variation was > 0.2 or the difference between ODs was greater than 0.2. We also checked the correlation of ODs across duplicate plates and re-ran both plates if r < 0.8. We included two sample free wells (50µl TBST) as blanks and two wells of positive controls on each plate. The positive control for the IgE assay was pooled serum from ewes trickle-infected with *T. circumcincta* and for the IgA and IgG assays was pooled plasma from normal healthy non-immunised domestic sheep. For subsequent analyses, the mean optical density ratio of each sample was taken according to this formula:

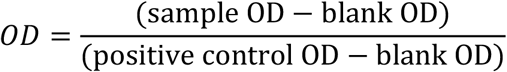

where the numerator was set to zero if the blank OD was greater than the sample OD in order to avoid negative values. Distributions of antibodies are shown in Figure S1. The number of samples that failed quality control per assay was 13 for IgA (7 lambs and 6 adults), 8 for IgE (6 lambs and 2 adults) and 27 for IgG (5 lambs and 22 adults). Correlations between antibody measures were modelled using linear regressions in R v3.4.3 (Figure S3).

### SNP data set

DNA was extracted from ear tissue or buffy coats using the Qiagen DNeasy blood and tissue kit according to the manufacturer’s protocol, except that a single final elution with 50ul AE buffer was used to give DNA at a concentration ≥ 50ng/ul. A total of 7,386 Soay sheep have been genotyped at 51,135 SNPs on the Illumina Ovine SNP50 BeadChip. Quality control was carried out using the check.marker function in GenABEL version 1.8-0 (56) using the following thresholds: SNP minor allele frequency (MAF) > 0.01, SNP locus genotyping success > 0.95, individual sheep genotyping success > 0.95, identity by state with another individual ≥ 0.95. Following quality control, 39,176 SNPs from 7,268 sheep remained. A further 189 sheep have been genotyped at 606,066 SNP loci on the Ovine Infinium HD SNP BeadChip and were subject to the same quality control thresholds as above (see (54) for individual selection criteria). All SNP locations were taken from their estimated positions on the sheep genome assembly Oar_v3.1 (GenBank assembly ID GCA_000298735.1; Jiang et al. 2014). Pedigree relationships between individuals were inferred using data from 438 SNP loci in the R package Sequoia v1.02 (58) and from field observations between mothers and their offspring born within the study area (see (53) for SNP selection criteria).

### Animal models

We modelled IgA, IgE and IgG levels in lambs and adults using a restricted maximum likelihood (REML) animal model approach (59) to determine the heritability of antibody levels in ASReml-R 3.0 (60) in R v3.4.3. We analysed lambs and adults separately due to a large difference observed in antibody levels (Figure S2) and due to the expected immaturity of the immune response in 4-month-old lambs (50). Using the above SNP dataset, a genomic relatedness matrix (GRM) at all autosomal markers was constructed for all genotyped individuals using GCTA 1.90.2 beta0 (61) to determine the variance attributed to additive genetic effects (i.e. the narrow-sense heritability, h^2^). Pedigree and GRM relatedness have been shown to be highly correlated in this system (62). The GRM was adjusted using the argument *--grm-adj 0*, which assumes that allele frequencies of causal and genotyped loci are similar. The fixed effect structure for the lamb-only models included sex and age in days as a linear covariate, while the random effects included the additive genetic component, maternal identity, birth year, ELISA plate number and ELISA run date. The fixed effect structure for the adult models included sex and age in years as a linear covariate, while the random effects included permanent environment (i.e. repeated measures within an individual) and capture year effects in addition to the random effects included in the lamb model. The proportion of the phenotypic variance explained by each random effect was estimated as the ratio of the relevant variance component to the sum of all variance components (i.e. the total phenotypic variance) as estimated by the animal model. The heritability of each measure was determined as the ratio of the additive genetic variance to the total phenotypic variance. The repeatability (i.e. the between-individual variation) of each measure in the adult and all age models was determined as the ratio of the sum of the additive genetic and permanent environment variance to the total phenotypic variance.

### Genome-wide association studies

Genome-wide association (GWA) was used to identify associations between individual single nucleotide polymorphisms (SNPs) and IgA, IgE and IgG levels in lambs and adults. This included SNPs on the X chromosome (N = 824) and those of unknown position (N = 313). For each trait and each class, a total of 39,176 individual animal models were run to determine the association with each SNP locus. Each model used the same fixed effect structures as above, with SNP genotype fitted as a two or three-level factor. To speed up computational time, the GRM was replaced with a relatedness matrix based on the pedigree (which is highly correlated with the GRM in this population (62)), and ELISA plate ID and run date were removed as random effects as they explained a very small proportion of the phenotypic variance (Figure 1). Models were run in ASReml-R 3.0 (60) in R v3.4.3. P-values were corrected for any additional unaccounted-for population structure by dividing them by the genomic control parameter λ (63) in cases where λ > 1, to reduce the incidence of false positives. λ was calculated as the median Wald test χ^2^_2_ divided by the median χ^2^_2_ expected from a null distribution. The significance threshold after multiple testing was determined using a linkage disequilibrium-based approach with a sliding window of 50 SNPs (outlined in (64)); for a false discovery rate of α = 0.05, the threshold P-value was set at 2.245×10^−6^ (54). Lamb IgE levels show strong right skew in their distribution (Figure S1), which can increase spurious associations at rare alleles present in individuals with large trait values. To mitigate against this, all zero trait values were removed (N = 394), and the response variable log_10_ transformed; this correction had a negligible effect on the variance component estimates.

**Figure 1:**
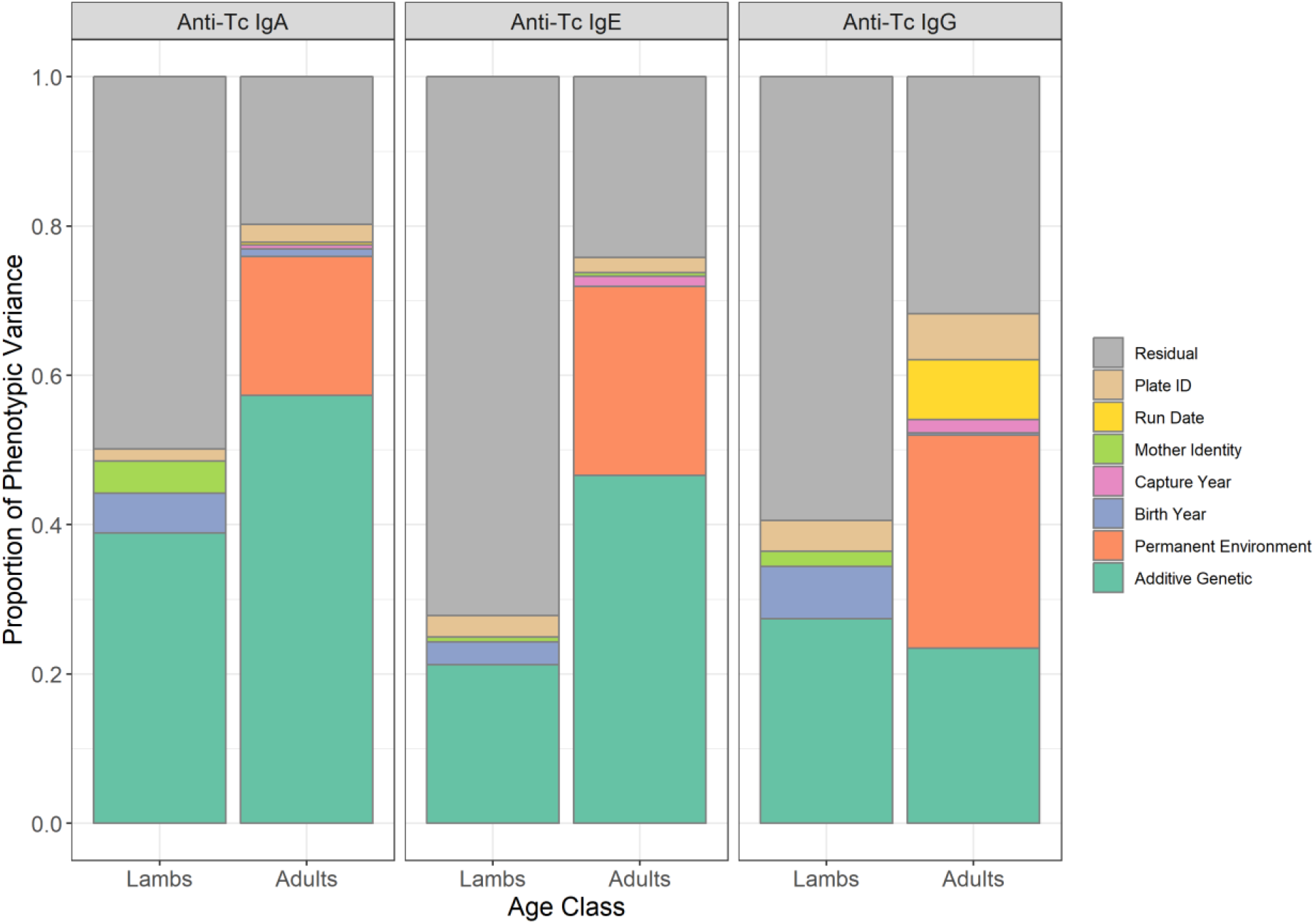
Proportion of phenotypic variance explained by random effects in animal models of anti-*T. circumcincta* IgA, IgE and IgG levels in lamb and adult Soay sheep. Data is provided in Table 1.

**Table 1.** Mean and variance estimates, and the proportion of variance explained for anti-*T. circumcincta* IgA, IgE and IgG levels measured in St. Kilda Soay sheep lambs and adults. Mean and V_OBS_ are the mean and variance of the raw data measures, N is the number of measures in N_IDS_ unique individuals. V_P_ is the phenotypic variance as a sum of all variance components as estimated by an animal model. The additive genetic effect (h^2^) indicates the narrow sense heritability of the trait. Non-significant estimates are indicated in grey text. Full results of all variance components are provided in Table S3. Figures in parentheses are standard errors.

| Trait | Age | $V_{OBS}$ | Mean | N | $N_{IDS}$ | $V_P$ | Proportion of $V_P$ explained | | | | | | | |
| --- | --- | --- | --- | --- | --- | --- | --- | --- | --- | --- | --- | --- | --- | --- |
| | | | | | | | Additive Genetic ($h^2$) | Permanent Environment | Birth Year | Capture Year | Mother Identity | Plate ID | Run Date | Residual |
| Anti-Tc IgA | Lambs | 0.2529 | 0.741 | 2030 | 2030 | 0.2483 | 0.3890 | NA | 0.0529 | NA | 0.0436 | 0.0157 | 0.0000 | 0.4989 |
|  |  |  |  |  |  | (0.0099) | (0.0372) | NA | (0.0202) | NA | (0.0188) | (0.0098) | (0.0000) | (0.0366) |
|  | Adults | 0.3051 | 1.507 | 3793 | 1321 | 0.3048 | 0.5732 | 0.1863 | 0.0102 | 0.0052 | 0.0000 | 0.0242 | 0.0032 | 0.1977 |
|  |  |  |  |  |  | (0.0134) | (0.0363) | (0.0303) | (0.0068) | (0.0035) | (0.0000) | (0.0075) | (0.0068) | (0.0101) |
| Anti-Tc IgE | Lambs | 0.0138 | 0.086 | 2035 | 2035 | 0.0135 | 0.2122 | NA | 0.0305 | NA | 0.0067 | 0.0288 | 0.0000 | 0.7219 |
|  |  |  |  |  |  | (0.0005) | (0.0334) | NA | (0.0153) | NA | (0.0174) | (0.0132) | (0.0000) | (0.0360) |
|  | Adults | 0.1835 | 0.733 | 3798 | 1321 | 0.1739 | 0.4662 | 0.2531 | 0.0000 | 0.0134 | 0.0047 | 0.0208 | 0.0000 | 0.2418 |
|  |  |  |  |  |  | (0.0071) | (0.0385) | (0.0368) | (0.0000) | (0.0057) | (0.0182) | (0.0055) | (0.0000) | (0.0117) |
| Anti-Tc IgG | Lambs | 0.0364 | 0.236 | 2032 | 2032 | 0.0354 | 0.2739 | NA | 0.0703 | NA | 0.0203 | 0.0411 | 0.0000 | 0.5944 |
|  |  |  |  |  |  | (0.0015) | (0.0344) | NA | (0.0266) | NA | (0.0184) | (0.0161) | (0.0000) | (0.0381) |
|  | Adults | 0.0462 | 0.630 | 3776 | 1319 | 0.0468 | 0.2347 | 0.2854 | 0.0027 | 0.0180 | 0.0000 | 0.0618 | 0.0803 | 0.3172 |
|  |  |  |  |  |  | (0.0021) | (0.0330) | (0.0296) | (0.0048) | (0.0101) | (0.0000) | (0.0172) | (0.0300) | (0.0159) |

### Variance explained by significantly associated regions

In regions of the genome where a SNP locus was significantly associated with an antibody measure, the proportion of phenotypic variation explained was modelled using a regional heritability approach (65). Briefly, a second GRM was constructed as above using SNP data from the most highly associated SNP in that region and the 9 SNP loci flanking that SNP on either side (i.e. 19 SNPs in total). This GRM was fitted as an additional random effect in the animal models and used to quantify the variance explained by variants within the associated region (see (54) for further details on the use of this method in Soay sheep).

### Imputation of SNP genotypes in associated regions

Further investigation of significant associations from GWAS was carried out using an imputation approach using data from individuals typed on the Ovine Infinium HD SNP BeadChip. SNP genotypes were extracted from the HD chip ±2Mb on either side of all significantly associated regions. Imputation was carried out for each region in AlphaImpute v1.9.8.4 (66,67) integrating pedigree information; parameter files for each region are included in the analysis code repository (see below). SNPs with an imputation success >95% were retained and associations between antibody levels and genotypes at each imputed SNP was calculated using the same animal model structures as outlined for the GWAS above.

### Gene and gene ontology annotation in associated regions

Gene annotations in significant regions were obtained from Ensembl (gene build ID Oar_v3.1.94). Gene ontology (GO) annotations for genes occurring within 1Mb of a significantly associated SNP were obtained from humans, mice, cattle and sheep gene builds using the function *getBM* in the R package biomaRt v2.34.2 (68). For genes where the gene name was not known, orthologous genes were identified using the biomaRt function *getLDS*. For all the genes and orthologues identified within these regions, the gene names, phenotype descriptions and GO terms were queried for all terms associated with immune function and antibodies (using the strings immun* and antibod*).

### Data availability

Data will be archived on a publicly accessible repository. All results and data underlying the figures in this manuscript are provided as Supplementary Material. All scripts for the analysis are provided at https://github.com/sejlab/Soay_Immune_GWAS.

## Results

### Phenotypic variation and animal models

August *T. circumcincta*-specific antibody levels of the three isotypes tested (IgA, IgG and IgE) were weakly positively correlated with each other, with slightly stronger correlations in lambs (adjusted R^2^ values from 0.078 to 0.175 in lambs, and from 0.005 to 0.012 in adults, P < 0.001; Figure S3, Table S1). Males had lower IgA levels as lambs and lower IgG levels as lambs and adults compared to females (Wald test P < 0.001, Figure S4, Table S2). All three antibody isotypes were positively associated with age in days in lambs and age in years in adults, except for adult IgG levels, which were negatively associated with age (Wald test P < 0.001, Figure S5, Table S2). Each antibody isotype was highly repeatable in adults (IgA = 0.76, IgE = 0.72, IgG = 0.52; Figure 1, Table 1), indicating some stability of individual antibody levels over the lifespan of animals. This is illustrated by a strong positive correlation between antibody measures taken in two consecutive years (IgA and IgE: slopes > 0.8, Adjusted R^2^ > 0.66; IgG: slope = 0.52, Adjusted R^2^ = 0.29; Figure S6, Table S4). All antibody measures were heritable in lambs and adults, with IgA levels showing the highest heritabilities (h^2^ = 0.39 & 0.57 for lambs and adults, respectively; Tables 1 & S3, Figure 1). Heritabilities in lambs and adults were 0.21 and 0.47 for IgE, and 0.29 and 0.23 for IgG, respectively (Tables 1 & S3, Figure 1). There was significant variation in antibody levels among birth years in lamb IgA and IgG measures, although the effect was small (≤ 7% of the phenotypic variance). There was a weakly significant maternal effect explaining < 4% of variation in lamb IgA levels (Figure 1, Tables 1 & S3). In adults, capture year explained < 1.5% of the phenotypic variance in all antibody measures. The full results of the animal models are provided in Tables S2 (fixed effect structures) and S3 (random effect structures).

### Genome-wide association studies

Genome-wide association and regional imputation studies identified several genomic regions associated with variation in anti-*T. circumcincta* IgA, IgE and IgG levels in lambs and adults (Figures 2 and S7, Tables 2 & S5). All test statistics were corrected using the genomic control parameter λ; this value was low for all 6 GWAS (λ < 1.074), indicating that population structure was adequately captured by fitting pedigree relatedness. Below, we discuss associations for each antibody separately, with summary information in Table 2. Full association results for genome-wide and imputed SNPs are provided in Tables S5 and S6, respectively. Information on genes and orthologues within associated regions are provided in Table S7 and immune GO terms associated with these genes are provided in Table S8.

**Table 2.** SNPs showing the strongest association with anti-*T. circumcincta* IgA, IgE and IgG levels in lambs and adults. The P-values provided in this table have not been corrected using genomic control to allow comparisons between directly genotyped and imputed SNPs. Asterisks next to the SNP name indicate that the most highly associated SNP was imputed from the high-density SNP chip. N 50K and N HD indicate how many SNPs were significantly associated with the trait in the same region for the 50K and HD SNP chips, respectively. A and B indicate the reference and alternate alleles at each SNP. MAF indicates the minor allele frequency (allele B); for imputed SNPs, this was calculated using the HD chip data only and not imputed genotypes. Effects AA, AB and BB are the effect sizes as calculated from the associated animal model. Full results including corrected P values are provided in Tables S5 and S6; gene and GO information is provided in Tables S7 & S8. Lamb IgE associations are given for the log_10_ of the antibody measures (see Methods).

| Trait | Age | Chr | Position | Highest Associated SNP | N 50K | N HD | P | A | B | MAF | Effect AA | Effect AB | Effect BB | Prop V<sub>A</sub> Explained | Closest Gene | Candidate Genes in Region |
| --- | --- | --- | --- | --- | --- | --- | --- | --- | --- | --- | --- | --- | --- | --- | --- | --- |
| Anti-Tc | Lambs | 18 | 68137231 | s03219.1 | 1 | 33 | 1.47e<sup>-07</sup> | A | G | 0.328 | 0.000 | 0.091 | 0.202 | 0.101 | *CDCA4* | *IGH* complex |
| IgA |  | 20 | 25196550 | oar3_OAR20_25196550* | 0 | 1 | 1.96e<sup>-06</sup> | A | G | 0.490 | 0.000 | -0.083 | -0.175 | 0.136 | *ELOVL5* | MHC II locus |
|  |  | 24 | 10616039 | oar3_OAR24_10616039* | 6 | 118 | 4.08e<sup>-39</sup> | A | G | 0.484 | 0.000 | -0.192 | -0.424 | 0.200 | *GSPT1* | *CIITA*, *CLEC16A* |
|  | Adults | 24 | 10858856 | oar3_OAR24_10858856* | 25 | 383 | 5.74e<sup>-71</sup> | A | G | 0.472 | 0.000 | -0.383 | -0.718 | 0.272 | *SNX29* | *CIITA*, *CLEC16A* |
| Anti-Tc | Lambs | 10 | 10333145 | oar3_OAR10_10333145* | 1 | 2 | 2.91e<sup>-07</sup> | G | A | 0.023 | 2.929 | 2.819 | 0.000 | 0.000 | *OLFM4* | *OLFM4* |
| IgE | Adults | 20 | 25781566 | OAR20_27259292.1* | 3 | 25 | 5.09e<sup>-09</sup> | A | G | 0.386 | 0.000 | -0.061 | -0.220 | 0.080 | *HLA-DRA* | MHC II locus |
| Anti-Tc | Lambs | 16 | 12632988 | oar3_OAR16_12632988* | 1 | 31 | 5.20e<sup>-07</sup> | A | G | 0.036 | 0.000 | 0.026 | 0.529 | 0.020 | *MAST4* | *CD180* |
| IgG |  | 20 | 30876754 | oar3_OAR20_30876754* | 1 | 6 | 2.44e<sup>-07</sup> | G | A | 0.211 | -0.104 | -0.072 | 0.000 | 0.077 | *TRIM38* | MHC I/II |

**Figure 2:**
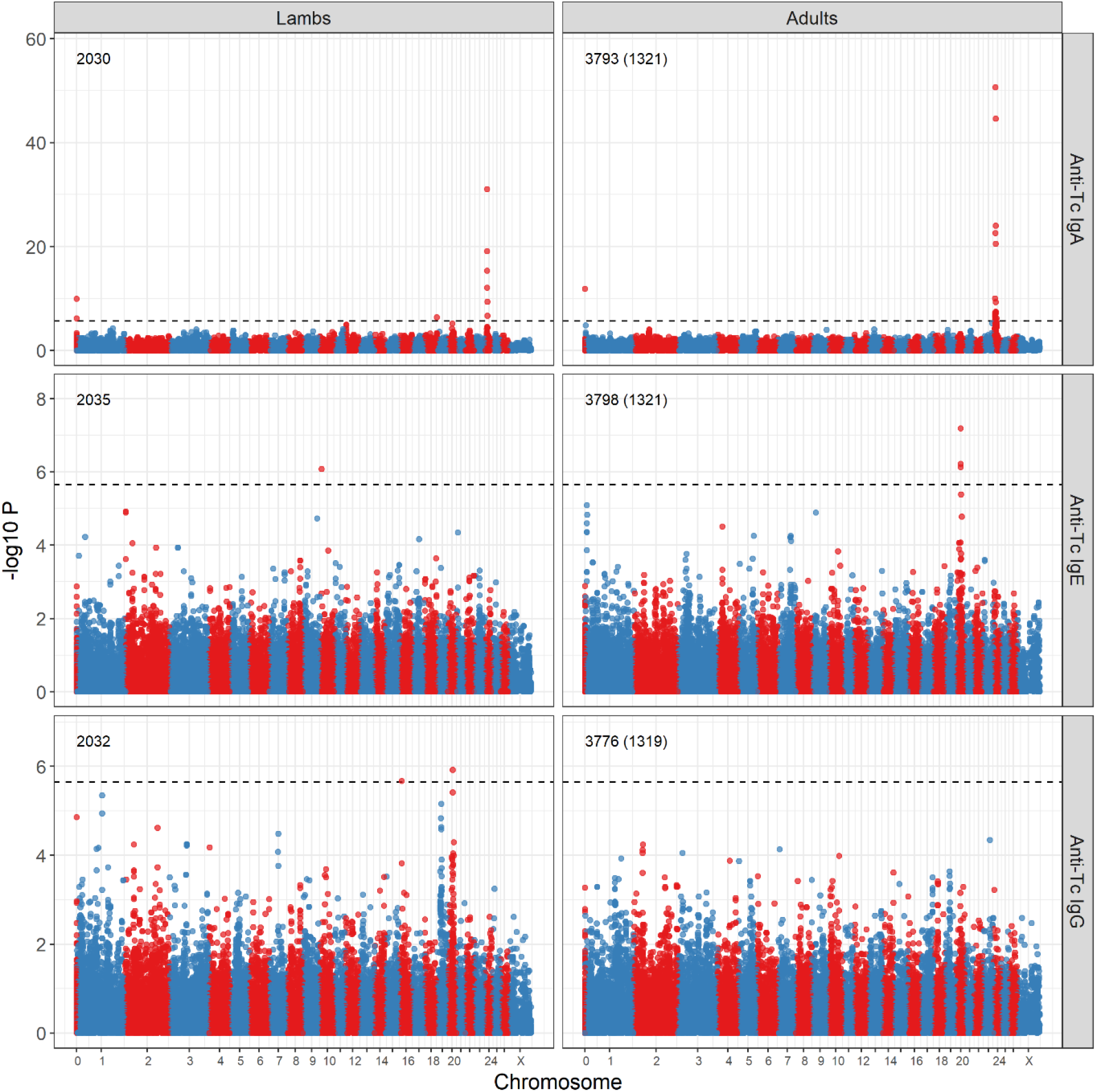
Genome-wide association of anti-*Teladorsagia circumcincta* IgA, IgE and IgG levels in lamb and adult Soay sheep with SNPs on the Ovine SNP50 BeadChip. Numbers indicate the number of measures and the number of unique individuals in parentheses. The dotted line indicates the genome-wide significance threshold equivalent to an experiment-wide threshold of P = 0.05. Points are colour-coded by chromosome. Positions are given relative to the sheep genome assembly Oar_v3.1. Underlying data, sample sizes and effect sizes are provided in Table S5. P-values were corrected with genomic control λ, and comparisons with those expected under a null distribution (i.e. P-P plots) are provided in Figure S7.

#### Anti-*T. circumcincta* IgA

There was a strong association between IgA levels and a region between 6.89 and 14.95 Mb on sheep chromosome 24, with the highest association observed at the SNP locus OAR24_12006191.1 in both lambs and adults (Wald test P = 1.01 ×10^−31^ and 2.23×10^−51^ in lambs and adults, respectively; Figures 2, 3 & S8; Tables 2 & S5). This SNP had an approximately additive effect on IgA levels in both lambs and adults (Table 2), with the region explaining 20.0% and 27.2% of the additive genetic variance in lambs and adults, respectively, equating to 7.8% and 15.3% of the phenotypic variance in lambs and adults, respectively. Associations at imputed SNPs in this region showed the strongest association at SNPs between 10.62Mb and 10.86Mb (maximum Wald test P = 4.08 ×10^−39^ and 5.74×10^−71^ in lambs and adults, respectively; Figure 3, Table S6), again with an additive effect on IgA levels (Table 2). This region corresponded to a novel gene (ENSOARG00000007156) orthologous to the protein coding gene Sorting Nexin 29 (*SNX29;* Figure 3, Table S7); GO terms indicated that this locus is associated with red blood cell phenotypes in humans and mice, including variation in haematocrit, erythrocyte cell number and circulating alkaline phosphate levels (International Mouse Phenotyping Consortium data (69); Table S8). Whilst this gene has no clear role in driving IgA levels, the associated SNPs were downstream of two candidate genes (Figure 3; Tables S7 & S8; distances of *~*1.021Mb and ~709Kb, respectively): the Class II Major Histocompatibility Complex Transactivator (*CIITA*), which is described as a “master control factor” for gene expression at the major histocompatibility complex (70,71); and C-Type Lectin Domain Containing 16A (*CLEC16A*), variants at which have been associated with common variable immunodeficiency disorder and IgA deficiency (72–74). An unmapped SNP was significantly associated with IgA levels in both lambs and adults (Figure 2, chromosome ‘0’); this locus was originally mapped to the same chromosome 24 region in version 2.0 of the sheep genome.

**Figure 3:**
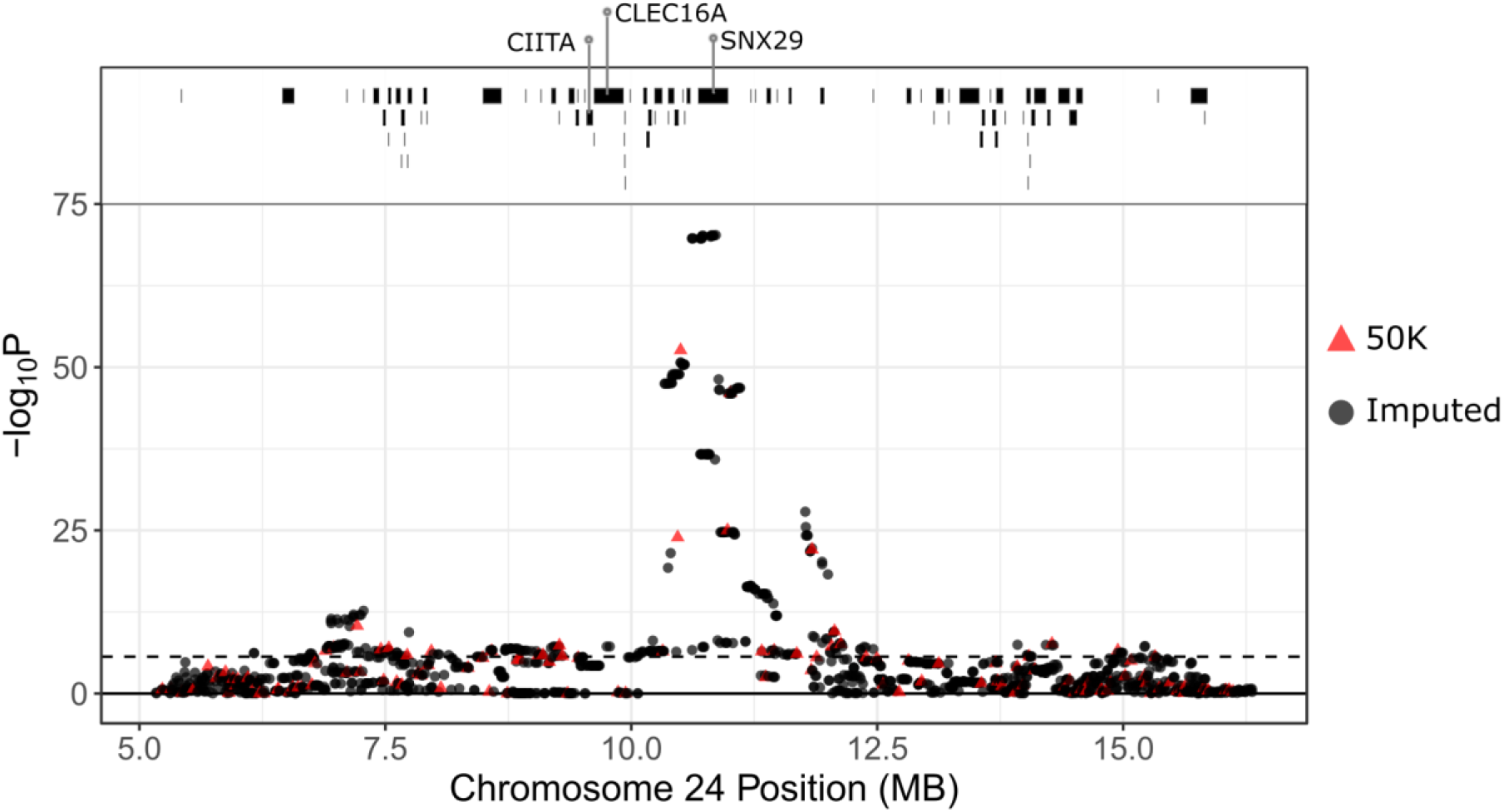
Local association of anti-*Teladorsagia circumcincta* IgA levels in adult Soay sheep with SNP50 and imputed SNP loci at the most highly associated region on chromosome 24. The dotted line indicates the genome-wide significance threshold equivalent to an experiment-wide threshold of P = 0.05. Points are colour-coded by their imputation status. Positions are given relative to the sheep genome assembly Oar_v3.1. Underlying data, sample sizes and effect sizes are provided in Table S6. Gene positions were obtained from Ensembl (gene build ID Oar_v3.1.94) and are provided in Table S7.

Lamb IgA levels showed a further association at a single common SNP at the distal end of chromosome 18 with an approximately additive effect on IgA levels (s03219.1, Wald test P = 4.34 ×10^−07^; Figures 2 & S8, Tables 2 & S5). No imputed loci passed the accuracy threshold in this region, meaning that fine mapping was not possible. Nevertheless, this SNP locus was located ~311kb to 454kb downstream of four novel genes (ENSOARG00000008862, ENSOARG00000008994, ENSOARG00000009143, ENSOARG00000009269) that are orthologous to various forms of immunoglobulin heavy constant alpha, epsilon, gamma and delta loci in humans (*IGHA*, *IGHE*, *IGHG* and *IGHD*, respectively; Tables S7 & S8). These loci code for constituent proteins of immunoglobulins and have GO terms associated with variation in IgA, IgE and IgG levels in mice (Table S8). A further association was observed at a single imputed SNP on chromosome 20, again with an approximately additive effect on IgA levels (oar3_OAR20_25196550, Wald test P = 1.96E-06; Figure S8; Tables 2 & S6) and ~ 157kb directly downstream of an orthologue of the MHC II locus *HLA-DQA1*.

#### Anti-*T. circumcincta* IgE

Lamb IgE levels were associated with a gene-poor region of chromosome 10, with the highest association observed at the imputed locus oar3_OAR10_10333145 (Figures 2 & S8, Table 2). The only protein-coding gene in this region, olfactomedin 4 (*OLFM4*), is associated with down-regulation of immune responses against bacterial infections in mice (75). However, given the low minor allele frequency of this locus, a lack of other associations at adjacent loci (Figure S8, Tables S5 & S6) and no contribution of the region to additive genetic variance (Table 2), we cannot rule out that association seen here is spurious and due to the sampling of rare alleles in individuals with extreme trait values.

Adult IgE levels were associated with a region from 25.8Mb to 27.5Mb on chromosome 20, with the highest association seen at the locus OAR20_27259292.1 for both the SNP50 and imputed SNP loci (Figures 2, Tables 2, S5 and S6). This SNP is directly upstream of the major histocompatibility complex (MHC) class II locus *HLA-DRA*, as well as the MHC class II loci *DQA* and *HLA-DQB1*; the wider region contains ~46 annotated genes with GO terms associated with immune function (Figure S8, Tables S7 & S8).

#### Anti-*T. circumcincta* IgG

Lamb IgG levels were significantly associated with a region from 29.6 – 30.9Mb on chromosome 20, with the highest association observed at the imputed locus oar3_OAR20_30876754 (Figures 2 & S8, Table 2). This region was ~4Mb from the region associated with IgE levels in adults and was close to protein coding regions orthologous to MHC Class I genes (Figure S8, Tables S7 & S8). A further association was observed on chromosome 16, corresponding to a region containing *CD180*, a gene associated with variation in IgG2b levels in mice (76) (Table S8), although the minor allele frequency of associated SNP is low (MAF = 0.036) and the association may again be partly driven by sampling effects (Table 2). There was no association between adult IgG levels and the SNPs genotyped in this study (Figure 2).

## Discussion

This study is one of the first to examine the genetic architecture of immune traits using a genome-wide association approach in a wild population. We have shown that anti-*Teladorsagia circumcincta* IgA, IgE and IgG levels in Soay sheep show substantial heritable variation underpinned by several genomic regions containing immune-associated genes. This suggests that antibody phenotypes have the potential to respond rapidly to selection, but also demonstrates that individual sheep develop distinct, temporally stable antibody phenotypes despite marked annual variation in exposure to nematode parasites, food availability and climate conditions (43,77,78). Below, we discuss the genetic architecture of these traits in more detail and how our findings inform the broader field of understanding the evolution and adaptive potential of immune traits in both domestic and natural populations.

### Temporal stability of antibody levels

We observed a large increase in anti-*Teladorsagia circumcincta* antibody levels between lambs (aged 4 months) and adults (aged >16 months). This was consistent with previous observations in this system and is probably due to the development of anti-helminth immunity with exposure over early life (24). In adults, antibody levels were stable within individuals, as indicated by high repeatabilities and strong temporal correlations of antibody measures between years (Figure S6, Table S4). This low intra-individual variation is notable given the temporally and spatially variable environment that individuals experience on St Kilda. The relatively small amount of variation explained by cohort, maternal and annual effects found here suggests that temporal variation in exposure to parasites, condition or early life effects had relatively little influence on antibody levels. It is also notable that repeatabilities for each antibody isotype were high despite different isotypes being only weakly correlated with one another, suggesting complex individualised immune phenotypes which are consistent over lifetimes. Our findings are consistent with the consensus emerging from human studies, which have also determined that variation in immune parameters is driven by high inter-individual and low intra-individual variation, indicative of stable immunological profiles of individuals (79,80). Whilst most intra-individual variation in this study was attributed to additive genetic effects, the permanent environment effects were substantial, accounting for 19%, 25% and 29% of the phenotypic variance in IgA, IgE and IgG, respectively. At present, the factors contributing to this variation remain unknown, but may be driven by consistent spatial differences in exposure or individual disease history, or due to complex interactions between nutritional state, exposure to other parasites and life history during early life.

### Heritable variation in antibody levels

Anti-*T. circumcincta* IgA, IgE and IgG levels were highly heritable in Soay sheep, ranging from 0.21 to 0.39 in lambs and from 0.23 to 0.57 in adults. These estimates are comparable to previous work estimating the pedigree heritability of an anti-*T. circumcincta* pan-isotype antibody measure (likely to be mainly comprised of IgG) in Soay sheep lambs (h^2^ = 0.30) and adults (h^2^ = 0.13 – 0.39) (26,47). In domestic sheep, similar heritability estimates have been obtained for anti-*T. circumcincta* IgE in Texel lambs (h^2^ = 0.39 and 0.50 against the third and fourth stage larvae, respectively (5)) and anti-*T. circumcincta* IgA in Scottish Blackface lambs (h^2^ = 0.56 against fourth stage larvae (6)). The observation that immune traits in Soay sheep and domestic breeds appear to have substantial heritable variation is interesting from an evolutionary perspective, as selection for reduced parasite load is likely to be strong, which in turn is predicted to reduce underlying genetic variation and hence the heritability of quantitative traits (81). In domestic sheep, anti-helminthic treatments may have relaxed the selection pressure on immune traits. Alternatively, the observed high heritabilities in both domestic and wild sheep may be in accordance with theory predicting that stabilising selection, rather than directional selection, is likely to be acting on immune traits (82), which in turn may lead to the maintenance of genetic variation at the underlying trait loci. In the Soay sheep, we have shown with the same dataset that there is little evidence for stabilising selection, with directional selection present for IgG in adults but not for other isotypes or age groups (50). It is notable that adult IgG, as well as being under the strongest directional selection, also has the lowest heritability compared to other isotypes and age groups (Figure 1, Table 1). This is consistent with the prediction that directional selection should erode heritable variation, whilst the high observed heritabilities in general are consistent with observations of weak or variable selection on these antibody measures (50). Nevertheless, a full understanding of the mechanisms maintaining this genetic variation will require examination of association between genotypes at significant loci with individual fitness, i.e. survival and reproductive success.

### Genetic variants associated with antibody levels

The strongest association observed in this study was between lamb and adult IgA levels and a region on chromosome 24 corresponding to the gene *SNX29*. This gene has no previous association with immune trait variation (see above) but occurs downstream of two candidate genes. The first, *CIITA*, is a master regulator of MHC class II gene expression; overexpression of *CIITA* in rats can induce transcription of MHC Class II genes in nearly all cell types (83) and *CIITA* knockout mice show impaired MHC Class II expression (84). Mutations in *CIITA* in humans are associated with bare lymphocyte syndrome type II, a severe primary immunodeficiency caused by the absence of MHC class II gene expression (85). In addition, a human GWAS study showed an association with variants at *CIITA* and levels of activated T cells (i.e., HLA DR+ T lymphocytes) and is in linkage disequilibrium with disease variants associated with ulcerative colitis (12). The second candidate, *CLEC16A*, is almost directly adjacent to *CIITA* and has been associated with IgA deficiency and common variable immunodeficiency disorder characterised by inadequate levels of multiple antibody isotypes (72–74). Further, *CLEC16A* knockdown mice have a reduced number of B cells and increased IgM levels compared with controls (73). Despite *CIITA* and *CLEC16A* being strong candidate genes for IgA expression *a priori*, they lie *~*1 & 0.7 Mb upstream from the GWAS peak, respectively (Figure 3). We cannot rule out that variants in protein-coding regions at *SNX29* and adjacent loci may drive IgA expression. However, a more plausible hypothesis is that the associated region contains cis-regulatory elements affecting the expression of *CIITA* and/or *CLEC16A*. Direct evidence of the precise cis-regulatory regions driving gene expression is scarce, but there is increasing evidence that genes can have multiple cis-regulatory regions driving expression (86), and that cis-regulatory regions can occur at distances of >1Mb from their target genes (see Orsolya & François 2013 and references therein).

The non-MHC variants identified in this study have not previously been associated with anti-*T. circumcincta* IgA, IgE or IgG levels in other sheep breeds investigated to date. A genome-wide association study in Scottish Blackface lambs failed to identify any SNPs associated with *T. circumcincta* IgA (37), while a study in Spanish Churra ewes found one genome-wide significant SNP on chromosome 12 (40). A quantitative trait locus (QTL) mapping study in Romney lambs found total IgE and anti-*Trichostrongylus colubriformis* IgG levels were each associated with a region on chromosome 23 (88). Together with our results, it appears that QTL for parasite-specific antibody traits have not been consistently observed between sheep breeds. This may be due to different loci associated with immune responses at different ages, differences in host-parasite exposure, inherent differences between breeds driven by different selective breeding histories, and/or genetic drift (26,40,89). Alternatively, there may be differences in the power to detect trait loci due to differences in patterns of linkage disequilibrium, effect sizes, sample sizes and/or analytical approaches between the studies. The loci identified in the current study may also be due to a genotype-by-environment effect that may only be manifested under natural conditions or could have been introduced with a historical admixture event with the Dunface breed (90). Investigation of candidate causal mutations in the current study will shed light on the mechanisms driving antibody levels within Soays, as well as their ubiquity and origin across different sheep breeds.

The identification of several large effect loci is in contrast with GWAS studies on body size and fitness-related traits in wild populations which have found few, if any, associations of SNPs with quantitative traits (27,53,91–95). This is because wild studies are subject to limitations related to sample size, environmental heterogeneity and marker density, which may fail to identify trait loci, over-estimate effect sizes and/or generate spurious associations (e.g. as stated above for observed associations at rare variants for lamb IgE and IgG on chromosomes 16 and 10, respectively) (96). We believe our overall findings are robust for the following reasons. This study has one of the highest sample sizes of any GWAS conducted in a wild system, with ~2,000 measures in lambs and ~3,800 measures in ~1300 unique adults, and sampling studies in this population suggest that causal variants contributing to heritable variation are adequately tagged by the Ovine SNP50 BeadChip (53,62). The extent of LD between genotyped SNP loci allowed successful imputation of high-density SNP loci in almost all significant regions of the genome, providing sufficient power to fine-map loci of large effect on immune phenotypes (53). We acknowledge that reduced LD in some regions (such as on chromosome 18 for lamb IgA) may mean that some regions of the genome are less able to tag heritable variation, potentially leading to reduced power to detect some trait loci. In addition, the Ovine SNP50 BeadChip has a low SNP density around the *DQA* and *DQB* loci in the MHC class II region, reducing power to detect associations (Figure S8b & S8f). Nevertheless: the quality of imputation was high within this region; other work has shown that there is no significant difference in patterns of LD and recombination rate compared to other locations within the genome (54); and traits were successfully mapped to the MHC region within the current study.

## Conclusion

This study provides evidence of a number of major effect loci and high additive genetic variation underlying complex immune traits in a wild population of Soay sheep, and provides a foundation for determining why genetic variation persists in immune traits by investigating associations with identified trait loci with individual fitness and genomic signatures of selection. The high heritability and repeatability of immune measures, as well as low correlations between them, suggests that strong targets for selection exist; a full understanding would require multivariate analysis to understand the constraints on immune phenotype evolution. Previous studies of immunity in the wild often focussed on specific immune regions (e.g. the MHC) and candidate genes encoding proteins of known immune function. Our study reveals the importance of using a genome-wide association, rather than candidate gene approach, for a clearer understanding of the genetic control of immune phenotypes. Overall, our study provides a rare example of multiple regions of large effect driving variation in immune phenotypes in the wild.

## Acknowledgements

We thank Ian Stevenson and all Soay sheep project members and volunteers for collection of data and samples. Phil Ellis, Camillo Bérénos and Hannah Lemon prepared DNA samples for Ovine SNP50 BeadChip genotyping and Jisca Huisman constructed the pedigree. Feedback from Jon Slate, Andrea Graham, Adam Hayward, Rick Maizels, Alastair Wilson and Amy Pedersen greatly improved this manuscript. This work has made extensive use of the Edinburgh Compute and Data Facility (http://www.ecdf.ed.ac.uk/). Permission to work on St Kilda is granted by The National Trust for Scotland, and logistical support was provided by QinetiQ, Eurest and Kilda Cruises. The Soay sheep project has been supported by grants from the UK Natural Environment Research Council. SNP genotyping was funded by a European Research Council Advanced Grant to JMP. DHN was supported by a Biotechnology and Biological Sciences Research Council David Phillips Fellowship. TNM is supported by the Scottish Government Rural Affairs, Food and the Environment (RAFE) Strategic Research Portfolio 2016-2021. AMS was supported by a Medical Research Council PhD Studentship. SEJ is supported by a Royal Society University Research Fellowship.

## Supplementary Information Description

**Figure S1.** Histograms of anti-*Teladorsagia circumcincta* IgA, IgE and IgG levels in lamb (left column) and adult (right column) Soay sheep.

**Figure S2.** Boxplots of anti-*Teladorsagia circumcincta* IgA, IgE and IgG levels with age and sex in Soay sheep.

**Figure S3.** Correlations between anti-*T. circumcincta* IgG, IgA, and IgE levels in lamb (A-C) and adult (D-F) Soay sheep. Model results are provided in Table S1.

**Figure S4.** Boxplots comparing anti-*T. circumcincta* IgG, IgA, and IgE levels between the sexes in lamb and adult Soay sheep.

**Figure S5.** Anti-*T. circumcincta* IgG, IgA, and IgE levels in lambs with age in days (left) and in adults with age in years (right). Animal model results are provided in Table S2.

**Figure S6.** Temporal correlations in anti-*Teladorsagia circumcincta* IgA, IgE and IgG levels in adult Soay sheep. Scatterplots of all raw data in adults for which there are two antibody measures in two consecutive years with a dashed line indicating a perfect 1:1 relationship and the solid line indicating the regression slope. Histograms show the frequency of the change in antibody levels for adults in consecutive years with a dashed line indicating no change.

**Figure S7:** Distribution of observed vs expected P-values under a null χ^2^ with 2 degrees of freedom for the GWAS of anti-*Teladorsagia circumcincta* IgA, IgE and IgG levels in lambs and adults. The dotted line indicates the genome-wide significance threshold, and the solid line indicates a 1:1 correspondence between the observed and expected values.

**Figure S8.** Local association of anti-*Teladorsagia circumcincta* IgA (a-d), IgE (e-f) and IgG (g-h) levels in lamb and adult Soay sheep with SNP50 and imputed SNP loci at the most highly associated regions. The dotted line indicates the genome-wide significance threshold equivalent to an experiment-wide threshold of P = 0.05. Points are colour-coded by their imputation status i.e. from the SNP50 chip (red points) or imputed from the Ovine HD chip (black triangles). Underlying data, sample sizes and effect sizes are provided in Table S6. Gene positions are shown in the grey panel at the top of each plot and were obtained from Ensembl (gene build ID Oar_v3.1.94) and are provided in Table S7. Genes coloured red have GO terms associated with immune traits (Table S8).

**Table S1:** Correlations between anti-*Teladorsagia circumcincta* antibody levels in lambs and adults. Slope, intercept, adjusted R^2^ and P-values are given for linear regressions.

**Table S2.** Fixed effects results from animal models of anti-*Teladorsagia circumcincta* IgA, IgE and IgG for lambs, and adults. Age is the age in days during the August catch for lambs, and age in years for adults. Wald statistics are given for the significance of each effect as included in the model.

**Table S3.** Random effects results from animal models of anti-*Teladorsagia circumcincta* IgA, IgE and IgG for lambs and adults. Wald statistics are given for the significance of each effect as included in the model. Sample sizes are provided in Table 1. Fixed effect structures and results are provided in Table S2.

**Table S4.** Temporal correlations in anti-*Teladorsagia circumcincta* IgA, IgE and IgG levels at time t and t+1 (in years) as shown in Figure S4. Results are from a linear regression with t+1 levels as the response variable.

**Table S5.** Full GWAS results for animal models of anti-*Teladorsagia circumcincta* IgA, IgE and IgG in lambs and adults, fitting SNP genotype as a factor. A and B indicate the reference and alternate allele at each SNP. CallRate is the genotyping success of the locus on the SNP50 BeadChip. MAF is the frequency of allele B (minor allele frequency). Wald P and Wald P Corrected are the association P-values before and after correction with genomic control λ, respectively. Significant indicates if the SNP was significantly associated with trait variation after correcting for multiple testing. Effect AA, AB and BB are the effect sizes from the animal model for each genotype relative to the model intercept.

**Table S6.** Full association results for animal models of anti-*Teladorsagia circumcincta* IgA, IgE and IgG in lambs and adults, fitting imputed SNP genotypes as a factor. SNP. Type indicates whether the SNP was imputed from the HD chip or from the SNP50 BeadChip (unknown genotypes are also imputed for the SNP50 BeadChip in this analysis meaning that results will not exactly match those of Table S5). A and B indicate the reference and alternate allele at each SNP. ImputeSuccess is the imputation success reported from the AlphaImpute analysis. MAF is the frequency of allele B (minor allele frequency). Wald P are the association P-values that have not been corrected for genomic control (see main text). Effect AA, AB and BB are the effect sizes from the animal model for each genotype relative to the model intercept.

**Table S7.** Gene information in regions significantly associated with anti-*Teladorsagia circumcincta* IgA, IgE and IgG in lambs and adults, obtained from the Ensembl Gene build Oar_v3.1.94. Start and stop indicate the gene start and stop positions. Strand indicates whether transcription occurs in the forward or reverse strand. Gene_id is the Ensembl identifier for the gene. Gene_name is the gene name associated with the gene_id. Gene_biotype indicates the type of gene (i.e. protein coding, RNA etc). Orthologue is the gene name of orthologues associated with the gene ID, with orthologue count giving the number of unique orthologues. Consensus locus is the gene name or likely gene name based on orthology.

**Table S8.** Gene Ontology information for loci (including orthologues) in Table S7 that are associated with immune and antibody phenotypes in humans (hsapiens), mice (mmusculus), cattle (btaurus) and sheep (oaries) obtained using biomaRt. Column names are as for Table S7, including the following: gene_id is the sheep gene ID; Species = species as previous; ensembl_gene_id is the gene ID within that Species; external_gene_name is the gene name for that species; description is the full gene name; phenotype_description is a description of phenotypes associated with the gene; go_id is the GO term identifier; name_1006 is the GO term name; definition_1006 is the GO term definition.

