## Supplementary figures and tables for "The genetic architecture of helminth-specific immune responses in a wild population of Soay sheep (*Ovis aries*)"

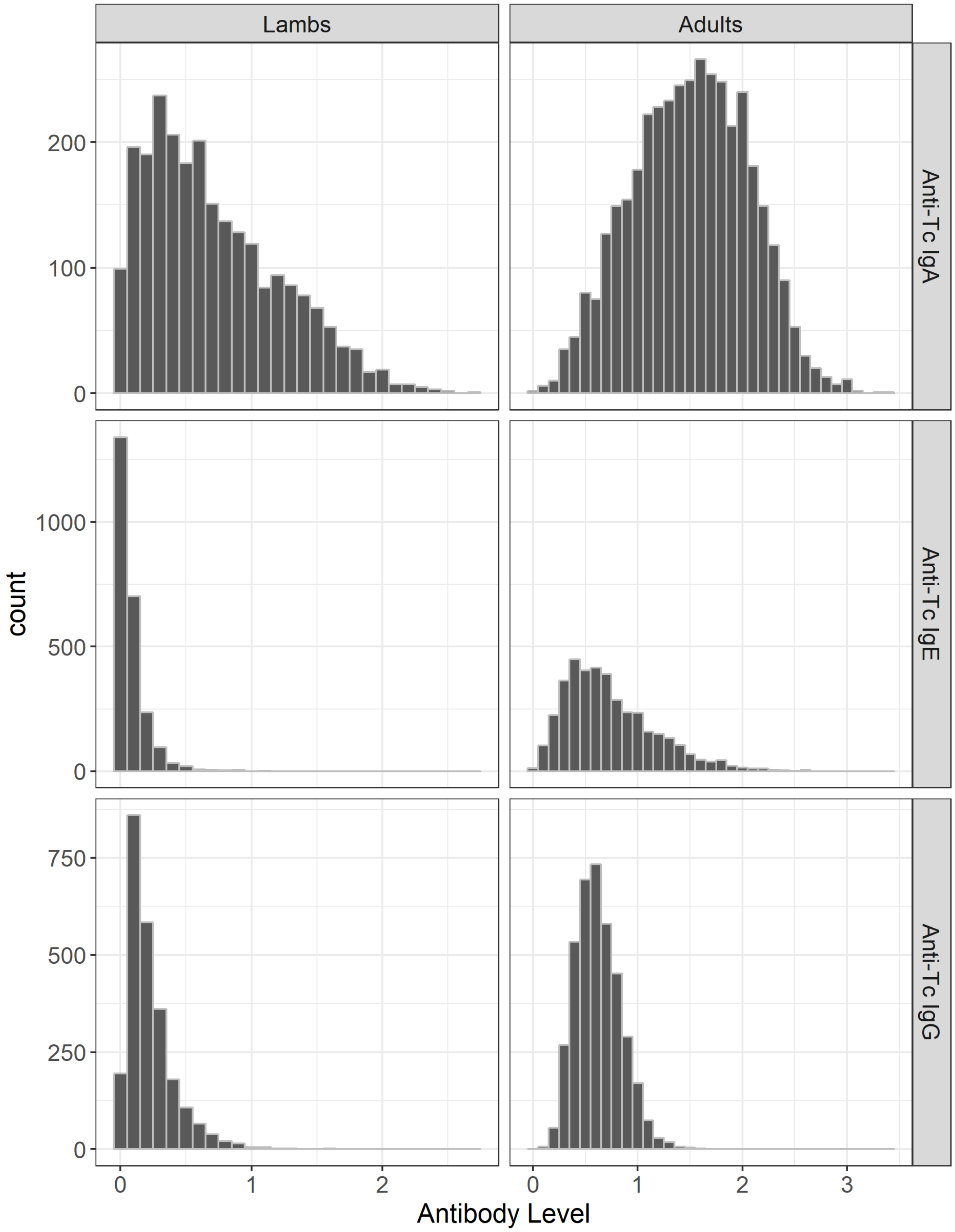


**Figure S1.** Histograms of anti-*Teladorsagia circumcincta* IgA, IgE and IgG levels in lamb (left column) and adult (right column) Soay sheep.

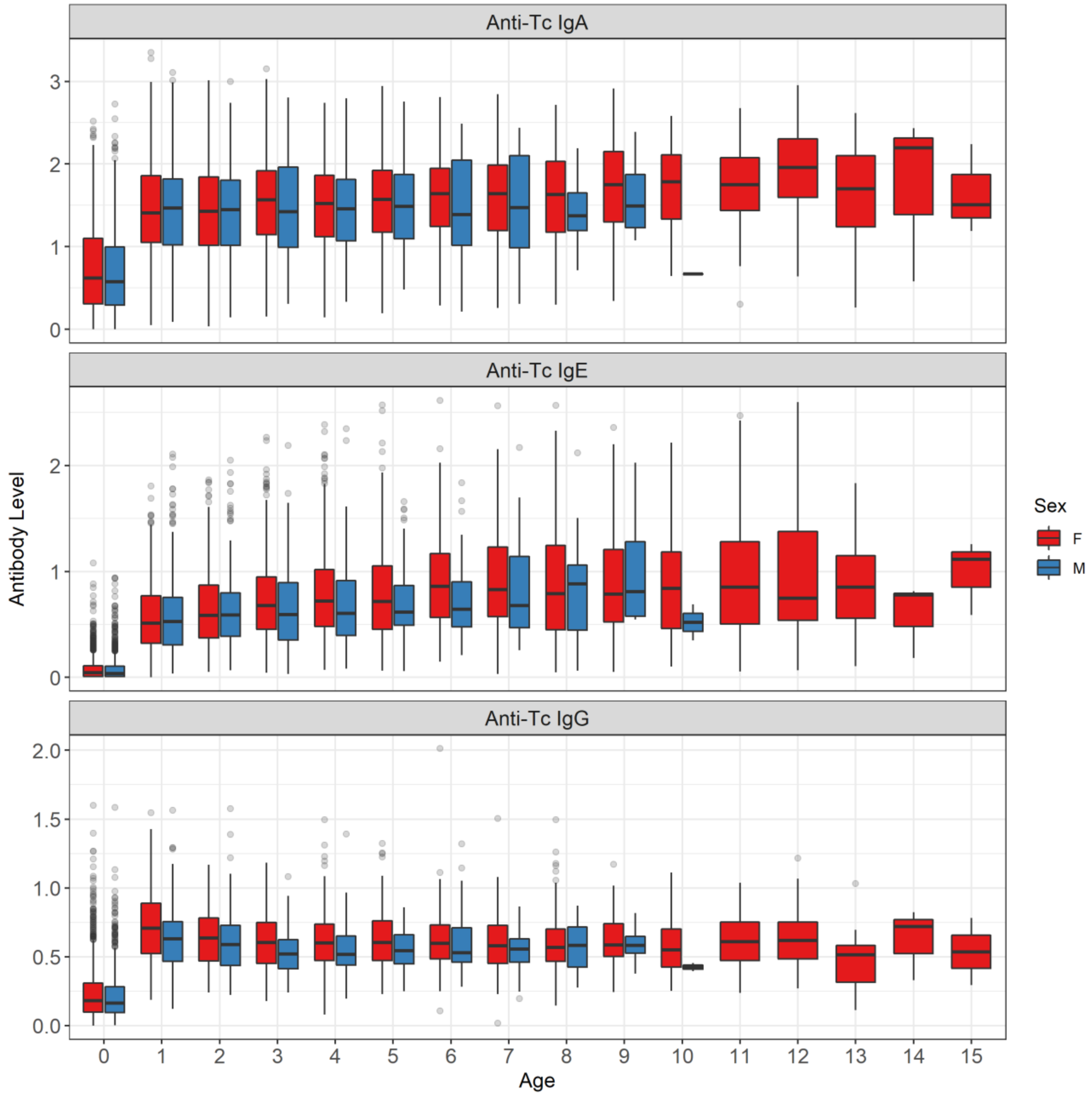


**Figure S2.** Boxplots of anti-*Teladorsagia circumcincta* IgA, IgE and IgG levels with age and sex in Soay sheep.

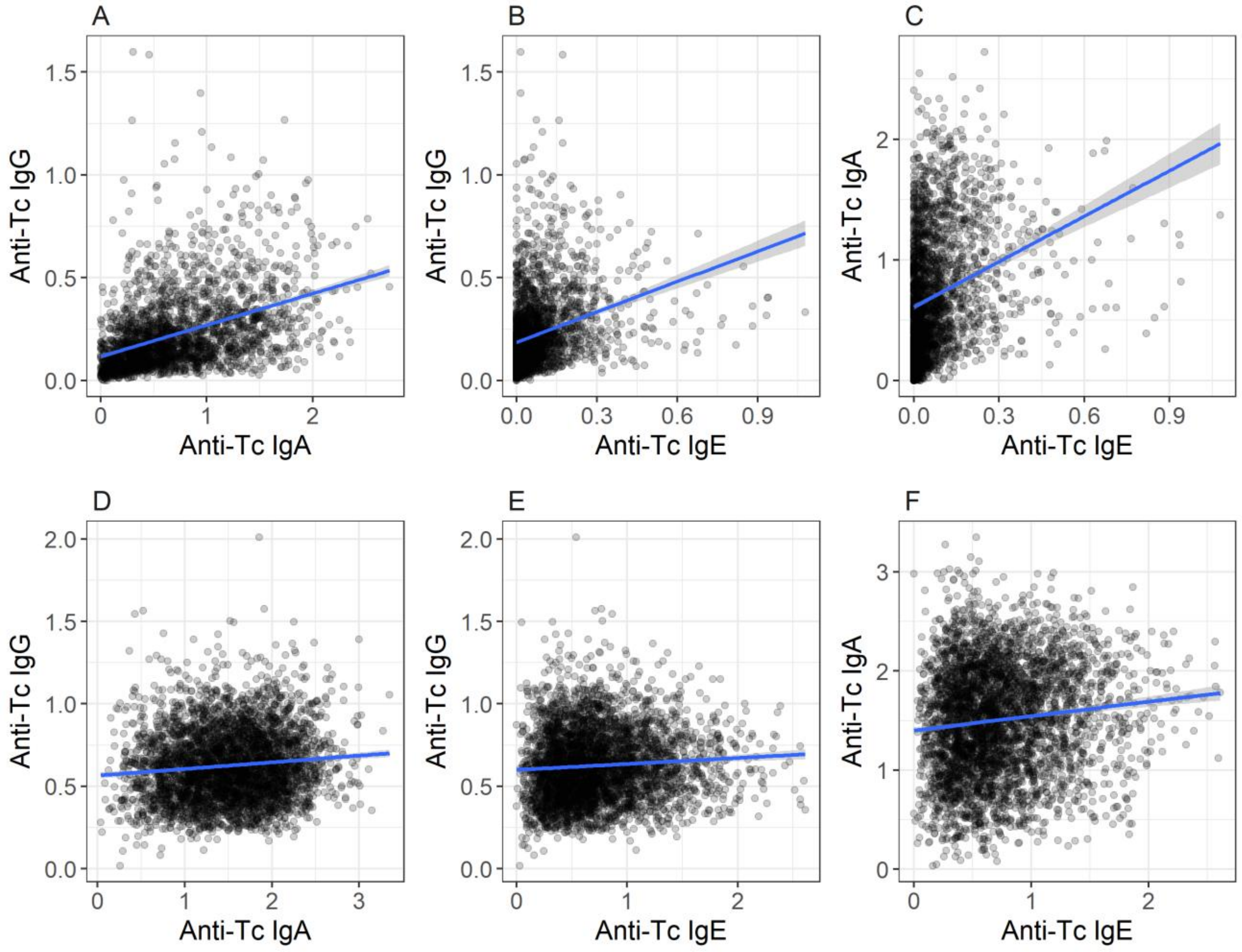


**Figure S3.** Correlations between anti-*T. circumcincta* IgG, IgA, and IgE levels in lamb (A-C) and adult (D-F) Soay sheep. Model results are provided in Table S1.

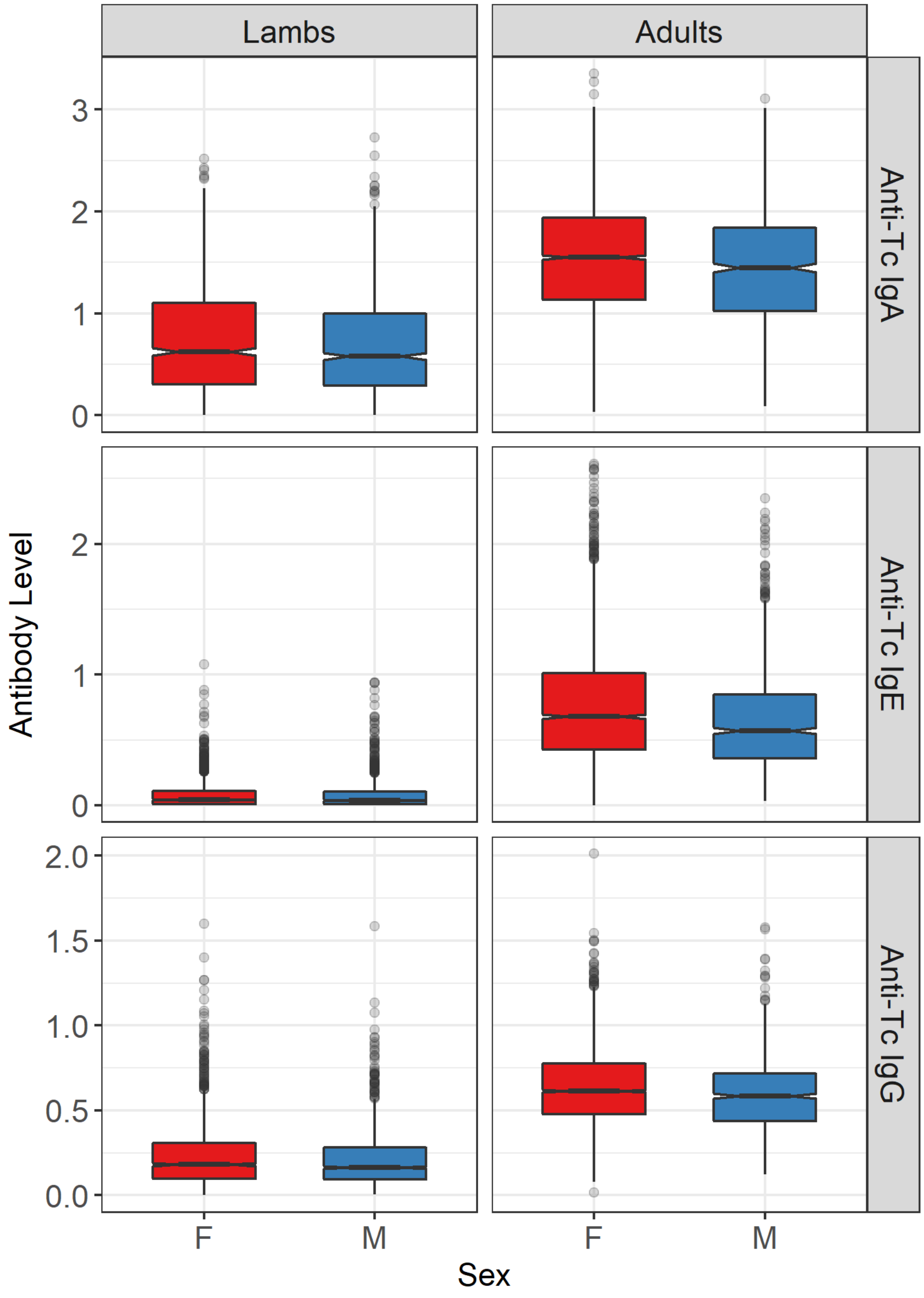


**Figure S4.** Boxplots comparing anti-*T. circumcincta* IgG, IgA, and IgE levels between the sexes in lamb and adult Soay sheep.

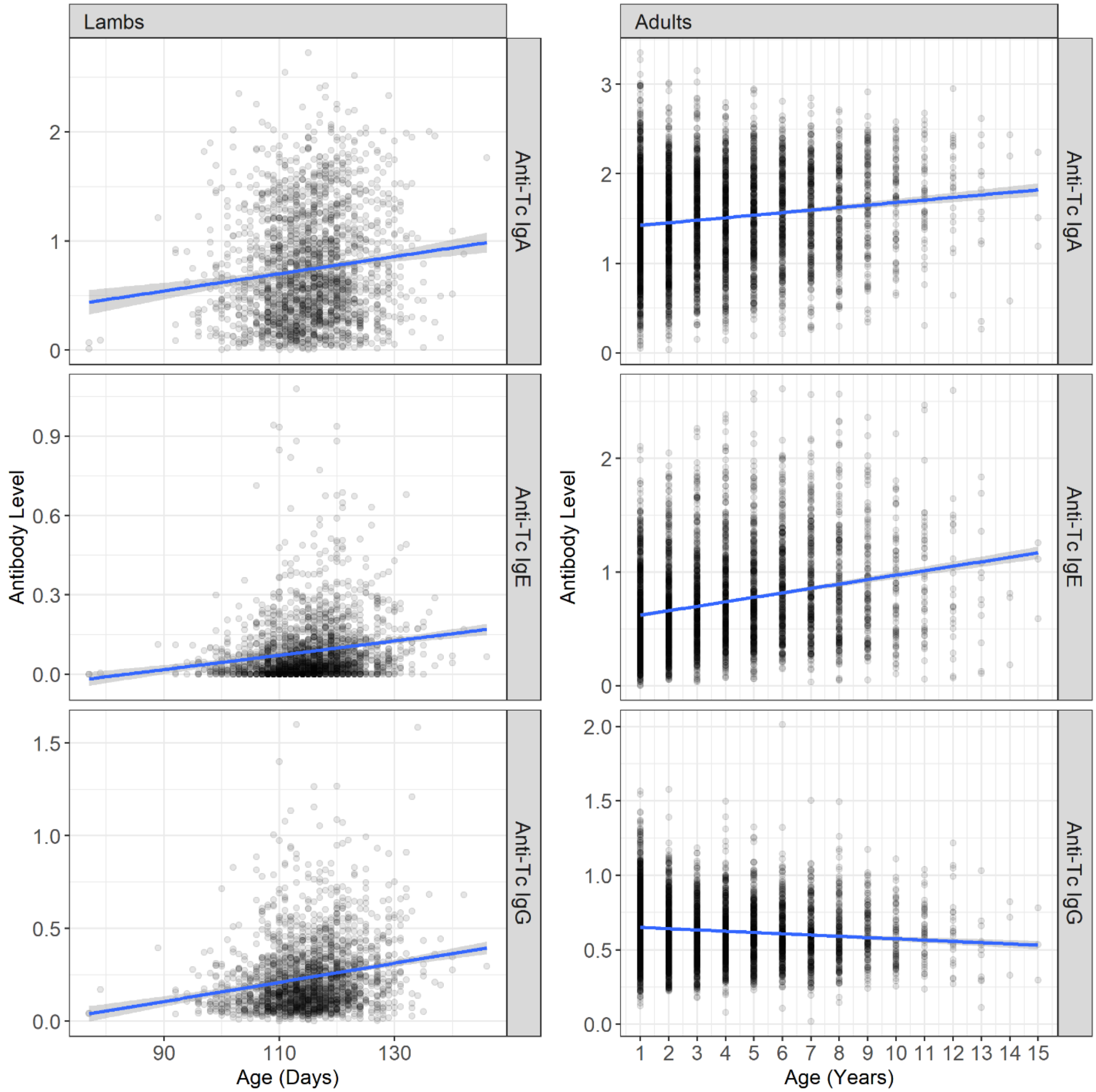


**Figure S5.** Anti-*T. circumcincta* IgG, IgA, and IgE levels in lambs with age in days (left) and in adults with age in years (right). Animal model results are provided in Table S2.

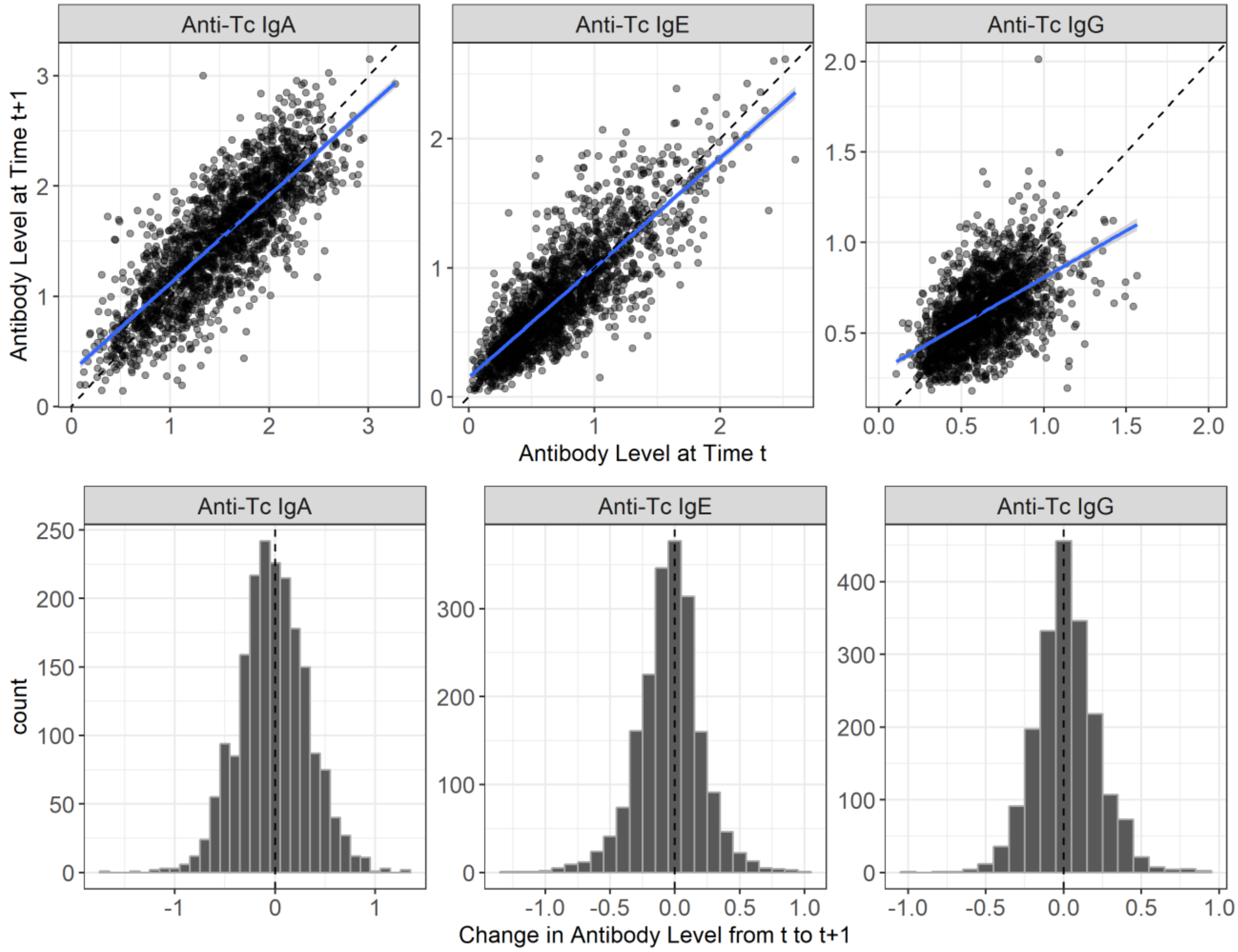


**Figure S6.** Temporal correlations in anti-*Teladorsagia circumcincta* IgA, IgE and IgG levels in adult Soay sheep. Scatterplots of all raw data in adults for which there are two antibody measures in two consecutive years with a dashed line indicating a perfect 1:1 relationship and the solid line indicating the regression slope. Histograms show the frequency of the change in antibody levels for adults in consecutive years with a dashed line indicating no change.

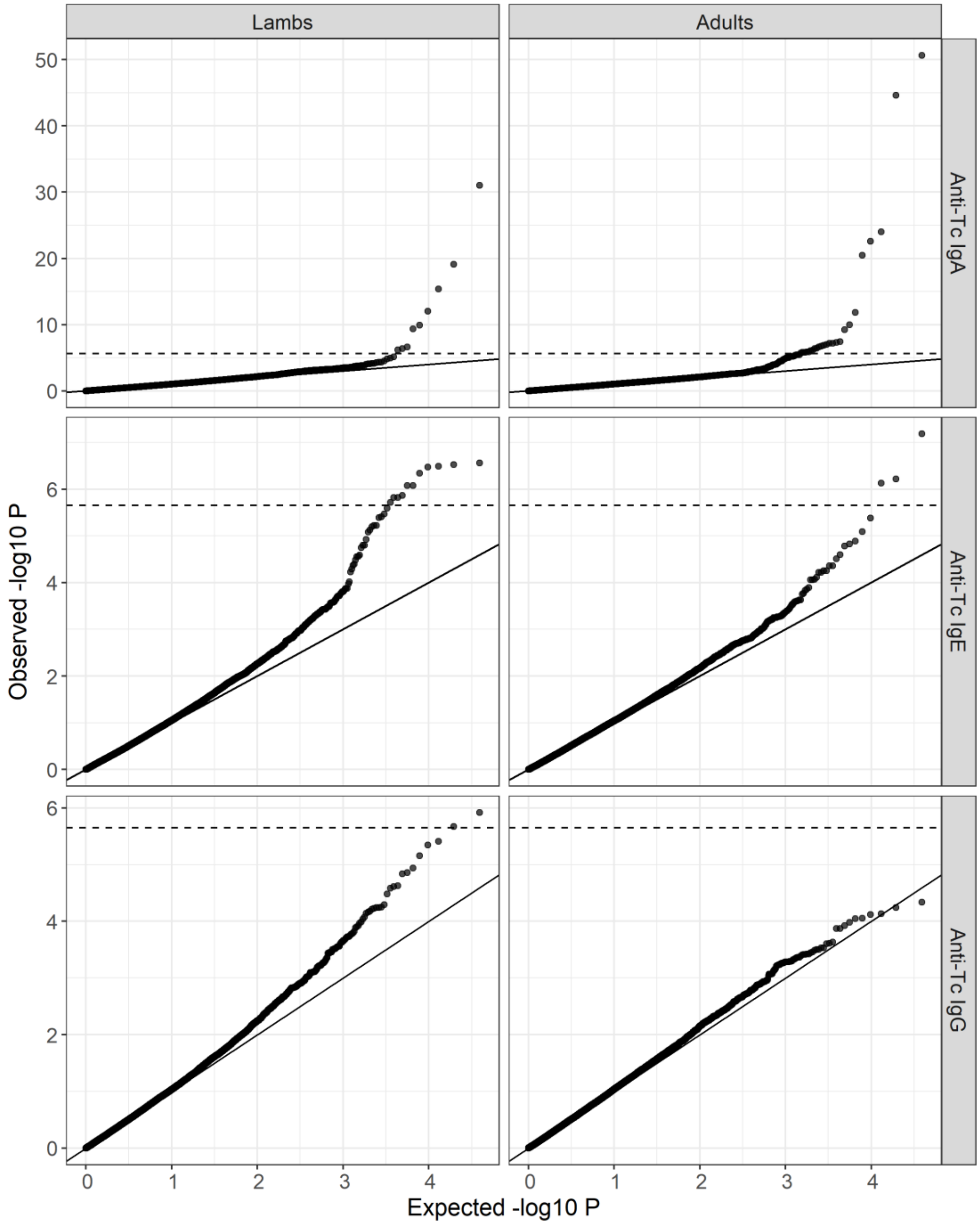


**Figure S7:** Distribution of observed vs expected P-values under a null $\chi^2$ with 2 degrees of freedom for the GWAS of anti-*Teladorsagia circumcincta* IgA, IgE and IgG levels in lambs and adults. The dotted line indicates the genome-wide significance threshold, and the solid line indicates a 1:1 correspondence between the observed and expected values.

**Figure S8.** Local association of anti-*Teladorsagia circumcincta* IgA (a-d), IgE (e-f) and IgG (g-h) levels in lamb and adult Soay sheep with SNP50 and imputed SNP loci at the most highly associated regions. The dotted line indicates the genome-wide significance threshold equivalent to an experiment-wide threshold of P = 0.05. Points are colour-coded by their imputation status i.e. from the SNP50 chip (red points) or imputed from the Ovine HD chip (black triangles). Underlying data, sample sizes and effect sizes are provided in Table S6. Gene positions are shown in the grey panel at the top of each plot and were obtained from Ensembl (gene build ID Oar_v3.1.94) and are provided in Table S7. Genes coloured red have GO terms associated with immune traits (Table S8).

a. Anti-Tc IgA: Lambs Chromosome 18

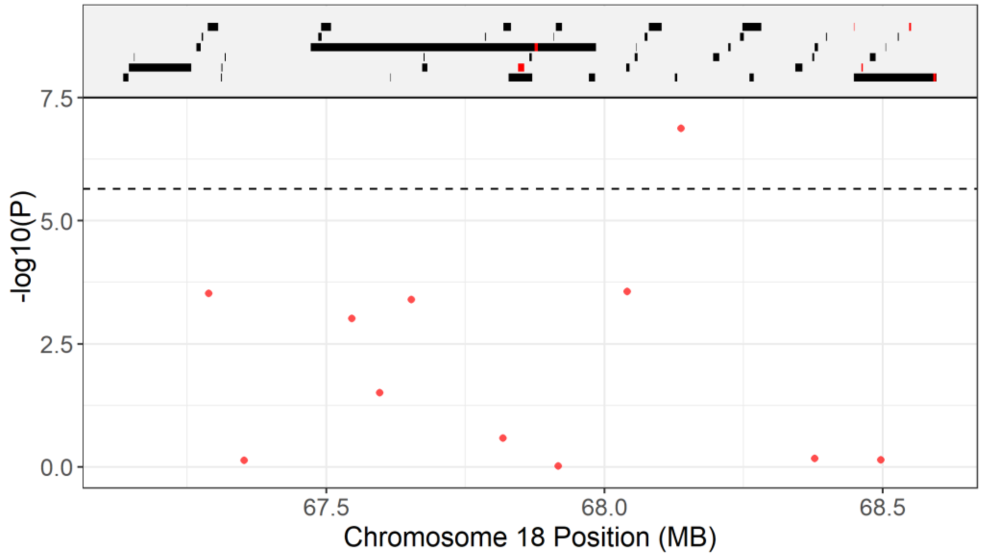


b. Anti-Tc IgA: Lambs Chromosome 20

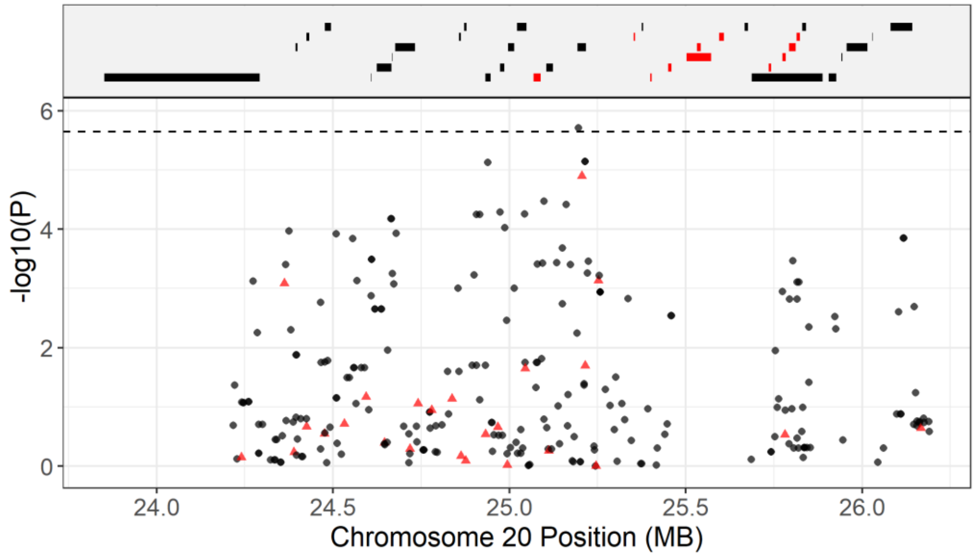

c. Anti-Tc IgA: Lambs Chromosome 24

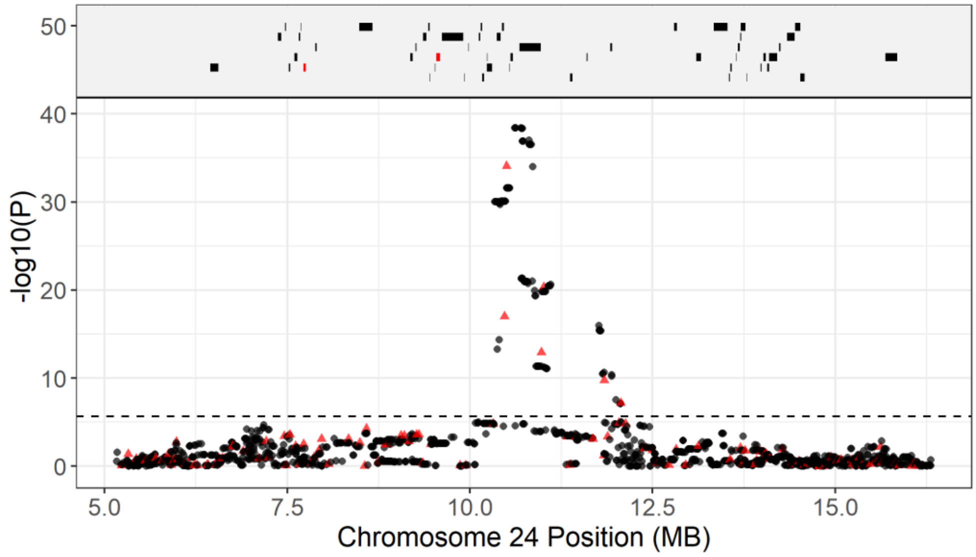


d. Anti-Tc IgA: Adults Chromosome 24

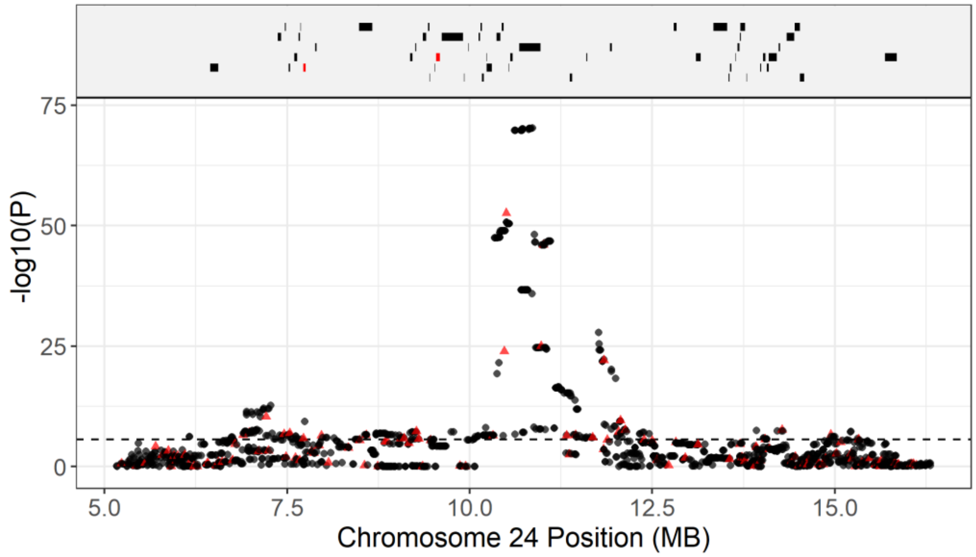

e. Anti-Tc IgE: Lambs Chromosome 10

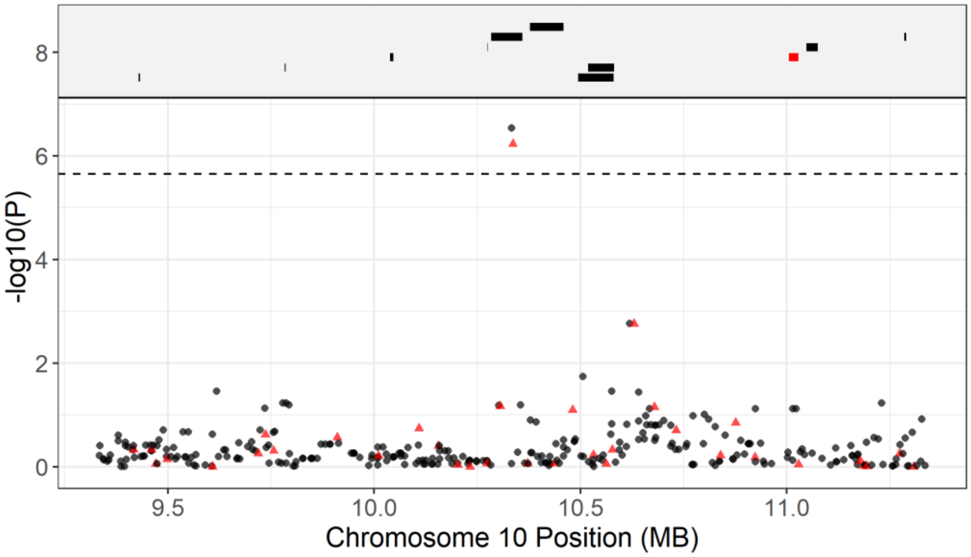


f. Anti-Tc IgE: Lambs Chromosome 20

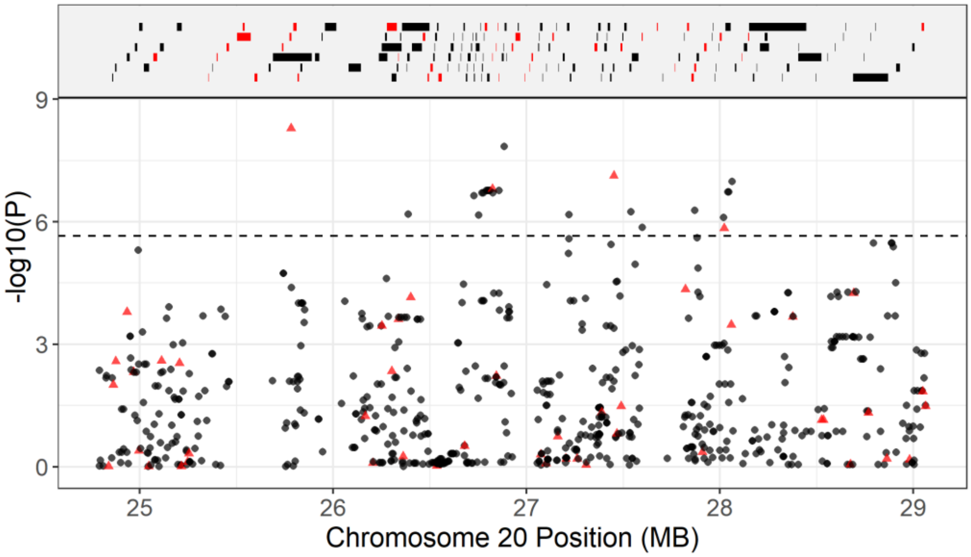

g. Anti-Tc IgG: Lambs Chromosome 16

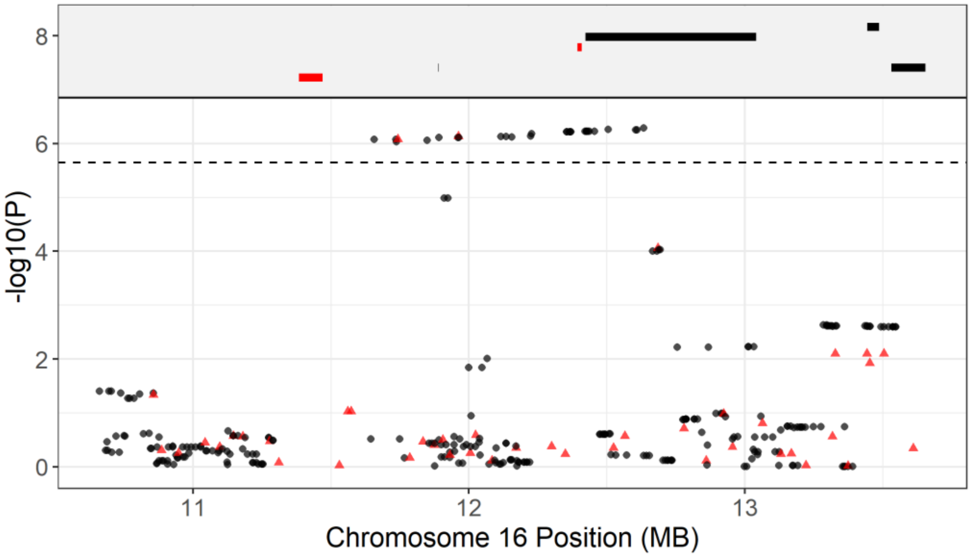


h. Anti-Tc IgG: Lambs Chromosome 20

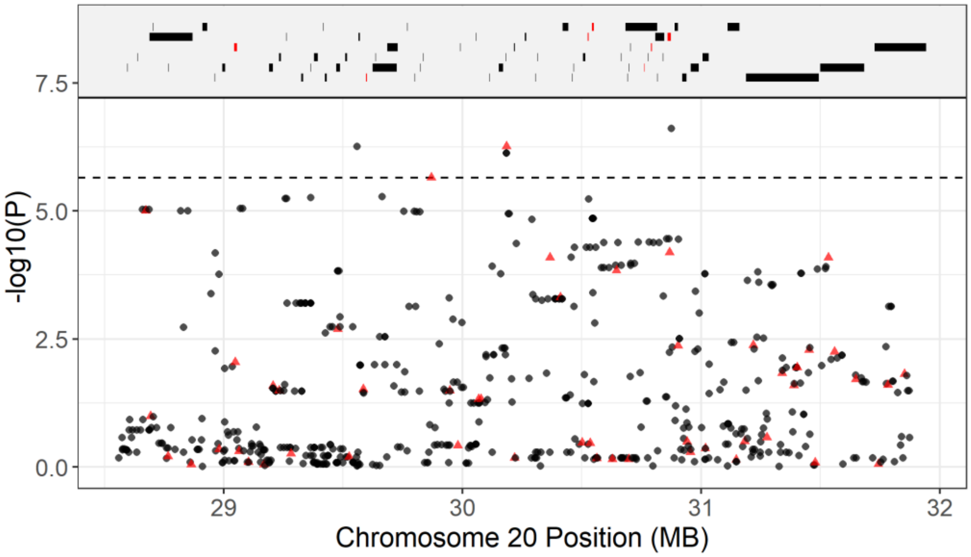

### Supplementary Tables

**Table S1:** Correlations between anti-*Teladorsagia circumcincta* antibody levels in lambs and adults. Slope, intercept, adjusted $R^2$ and P-values are given for linear regressions.

| Measure X | Measure Y | age | slope | intercept | Adj.R2 | P |
|---|---|---|---|---|---|---|
| Anti-Tc IgA | Anti-Tc IgG | Lambs | 0.153 | 0.117 | 0.175 | 5.29E-104 |
| Anti-Tc IgE | Anti-Tc IgG | Lambs | 0.492 | 0.186 | 0.09 | 4.15E-52 |
| Anti-Tc IgE | Anti-Tc IgA | Lambs | 1.256 | 0.61 | 0.078 | 2.56E-45 |
| Anti-Tc IgA | Anti-Tc IgG | Adults | 0.04 | 0.565 | 0.01 | 1.16E-10 |
| Anti-Tc IgE | Anti-Tc IgG | Adults | 0.036 | 0.599 | 0.005 | 6.47E-06 |
| Anti-Tc IgE | Anti-Tc IgA | Adults | 0.145 | 1.4 | 0.012 | 1.28E-12 |

**Table S2.** Fixed effects results from animal models of anti-*Teladorsagia circumcincta* IgA, IgE and IgG for lambs, and adults. Age is the age in days during the August catch for lambs, and age in years for adults. Wald statistics are given for the significance of each effect as included in the model.

| Trait | Age | Effect | Effect Level | Solution | Standard Error | Z.Ratio | Wald Statistic | Wald DF | P |
|---|---|---|---|---|---|---|---|---|---|
| Anti-Tc IgA | Lambs | (Intercept) | | -0.281 | 0.199 | -1.415 | 770.430 | 1 | 0 |
| | | Age | | 0.009 | 0.002 | 5.387 | 29.024 | 1 | 7.15E-08 |
| | | Sex | F | 0.000 | NA | NA | 11.742 | 1 | 6.11 E -4 |
| | | Sex | M | -0.065 | 0.019 | -3.341 | | | |
| | Adults | (Intercept) | | 1.408 | 0.027 | 52.818 | 4383.562 | 1 | 0 |
| | | Age | | 0.020 | 0.003 | 6.433 | 41.379 | 1 | 1.25E-10 |
| | | Sex | F | 0.000 | NA | NA | 0.502 | 1 | 0.479 |
| | | Sex | M | -0.010 | 0.025 | -0.408 | | | |
| Anti-Tc IgE | Lambs | (Intercept) | | -0.244 | 0.048 | -5.112 | 249.859 | 1 | 0 |
| | | Age | | 0.003 | 0.000 | 7.033 | 49.467 | 1 | 2.02E-12 |
| | | Sex | F | 0.000 | NA | NA | 0.300 | 1 | 0.584 |
| | | Sex | M | -0.002 | 0.005 | -0.431 | | | |
| | Adults | (Intercept) | | 0.613 | 0.019 | 31.622 | 1974.675 | 1 | 0 |
| | | Age | | 0.029 | 0.003 | 11.756 | 138.196 | 1 | 0 |
| | | Sex | F | 0.000 | NA | NA | 2.662 | 1 | 0.103 |
| | | Sex | M | -0.021 | 0.020 | -1.031 | | | |
| Anti-Tc IgG | Lambs | (Intercept) | | -0.459 | 0.078 | -5.905 | 409.237 | 1 | 0 |
| | | Age | | 0.006 | 0.001 | 9.325 | 86.950 | 1 | 0 |
| | | Sex | F | 0.000 | NA | NA | 15.708 | 1 | 7.39E-05 |
| | | Sex | M | -0.029 | 0.007 | -3.837 | | | |
| | Adults | (Intercept) | | 0.689 | 0.016 | 41.915 | 1729.272 | 1 | 0 |
| | | Age | | -0.015 | 0.002 | -8.845 | 78.234 | 1 | 0 |
| | | Sex | F | 0.000 | NA | NA | 36.348 | 1 | 1.65E-09 |
| | | Sex | M | -0.066 | 0.010 | -6.502 | | | |

**Table S3.** Random effects results from animal models of anti-*Teladorsagia circumcincta* IgA, IgE and IgG for lambs and adults. Wald statistics are given for the significance of each effect as included in the model. Sample sizes are provided in Table 1. Fixed effect structures and results are provided in Table S2.

| Trait | Age | Random Effect | Variance Component | Standard Error | Z Ratio | Effect (Proportion of Variance) | Standard Error |
|---|---|---|---|---|---|---|---|
| Anti-Tc IgA | Lambs | Birth Year | 0.0131 | 0.0052 | 2.5088 | 0.0529 | 0.0202 |
| | | Run Date | 0.0000 | 0.0000 | 16.2020 | 0.0000 | 0.0000 |
| | | Plate ID | 0.0039 | 0.0025 | 1.5831 | 0.0157 | 0.0098 |
| | | Mother Identity | 0.0108 | 0.0047 | 2.3049 | 0.0436 | 0.0188 |
| | | Additive Genetic | 0.0966 | 0.0110 | 8.7613 | 0.3890 | 0.0372 |
| | | Residual | 0.1239 | 0.0076 | 16.2020 | 0.4989 | 0.0366 |
| | Adults | Capture Year | 0.0016 | 0.0011 | 1.4760 | 0.0052 | 0.0035 |
| | | Birth Year | 0.0031 | 0.0021 | 1.4831 | 0.0102 | 0.0068 |
| | | Run Date | 0.0010 | 0.0021 | 0.4773 | 0.0032 | 0.0068 |
| | | Plate ID | 0.0074 | 0.0023 | 3.2042 | 0.0242 | 0.0075 |
| | | Mother Identity | 0.0000 | 0.0000 | 34.6559 | 0.0000 | 0.0000 |
| | | Additive Genetic | 0.1747 | 0.0170 | 10.2746 | 0.5732 | 0.0363 |
| | | Perm Environment | 0.0568 | 0.0083 | 6.8239 | 0.1863 | 0.0303 |
| | | Residual | 0.0603 | 0.0017 | 34.6559 | 0.1977 | 0.0101 |
| Anti-Tc IgE | Lambs | Birth Year | 0.0004 | 0.0002 | 1.9569 | 0.0305 | 0.0153 |
| | | Run Date | 0.0000 | 0.0000 | 20.5645 | 0.0000 | 0.0000 |
| | | Plate ID | 0.0004 | 0.0002 | 2.1460 | 0.0288 | 0.0132 |
| | | Mother Identity | 0.0001 | 0.0002 | 0.3847 | 0.0067 | 0.0174 |
| | | Additive Genetic | 0.0029 | 0.0005 | 5.9041 | 0.2122 | 0.0334 |
| | | Residual | 0.0097 | 0.0005 | 20.5645 | 0.7219 | 0.0360 |
| | Adults | Capture Year | 0.0023 | 0.0010 | 2.3497 | 0.0134 | 0.0057 |
| | | Birth Year | 0.0000 | 0.0000 | 34.8017 | 0.0000 | 0.0000 |
| | | Run Date | 0.0000 | 0.0000 | 34.8017 | 0.0000 | 0.0000 |
| | | Plate ID | 0.0036 | 0.0010 | 3.7558 | 0.0208 | 0.0055 |
| | | Mother Identity | 0.0008 | 0.0032 | 0.2559 | 0.0047 | 0.0182 |
| | | Additive Genetic | 0.0811 | 0.0091 | 8.9316 | 0.4662 | 0.0385 |
| | | Perm Environment | 0.0440 | 0.0059 | 7.4223 | 0.2531 | 0.0368 |
| | | Residual | 0.0420 | 0.0012 | 34.8017 | 0.2418 | 0.0117 |
| Anti-Tc IgG | Lambs | Birth Year | 0.0025 | 0.0010 | 2.4917 | 0.0703 | 0.0266 |
| | | Run Date | 0.0000 | 0.0000 | 18.4285 | 0.0000 | 0.0000 |
| | | Plate ID | 0.0015 | 0.0006 | 2.5077 | 0.0411 | 0.0161 |
| | | Mother Identity | 0.0007 | 0.0007 | 1.0990 | 0.0203 | 0.0184 |
| | | Additive Genetic | 0.0097 | 0.0013 | 7.2460 | 0.2739 | 0.0344 |
| | | Residual | 0.0210 | 0.0011 | 18.4285 | 0.5944 | 0.0381 |
| | Adults | Capture Year | 0.0008 | 0.0005 | 1.7634 | 0.0180 | 0.0101 |
| | | Birth Year | 0.0001 | 0.0002 | 0.5680 | 0.0027 | 0.0048 |
| | | Run Date | 0.0038 | 0.0015 | 2.5122 | 0.0803 | 0.0300 |
| | | Plate ID | 0.0029 | 0.0008 | 3.5223 | 0.0618 | 0.0172 |
| | | Mother Identity | 0.0000 | 0.0000 | 34.6355 | 0.0000 | 0.0000 |
| | | Additive Genetic | 0.0110 | 0.0017 | 6.4888 | 0.2347 | 0.0330 |
| | | Perm Environment | 0.0134 | 0.0013 | 10.0241 | 0.2854 | 0.0296 |
| | | Residual | 0.0149 | 0.0004 | 34.6355 | 0.3172 | 0.0159 |

**Table S4.** Temporal correlations in anti-*Teladorsagia circumcincta* IgA, IgE and IgG levels at time t and t+1 (in years) as shown in Figure S4. Results are from a linear regression with t+1 levels as the response variable.

| Model | Slope | Intercept | Adjusted R2 | P |
|---|---|---|---|---|
| Anti-Tc IgA | 0.801 | 0.318 | 0.656 | 0.00E+00 |
| Anti-Tc IgE | 0.848 | 0.154 | 0.69 | 0.00E+00 |
| Anti-Tc IgG | 0.519 | 0.287 | 0.293 | 2.30E-146 |

**Table S5.** Full GWAS results for animal models of anti-*Teladorsagia circumcincta* IgA, IgE and IgG in lambs and adults, fitting SNP genotype as a factor. A and B indicate the reference and alternate allele at each SNP. CallRate is the genotyping success of the locus on the SNP50 BeadChip. MAF is the frequency of allele B (minor allele frequency). Wald P and Wald P Corrected are the association P-values before and after correction with genomic control $\lambda$, respectively. Significant indicates if the SNP was significantly associated with trait variation after correcting for multiple testing. Effect AA, AB and BB are the effect sizes from the animal model for each genotype relative to the model intercept.

[ See CSV file Table_S8_Immune_GO_Terms_for_Significant_Regions ]
